# Insertions in universal proteins confirm that *Nanohaloarchaea* belong to DPANN*-Archaea* and suggest several new major archaeal clades

**DOI:** 10.64898/2026.06.17.729505

**Authors:** Patrick Forterre, Emmanuelle Schmitt, Violette Da Cunha

## Abstract

The phylogenetic position of *Nanohaloarchaea* has been debated, these nanosized archaea being alternatively proposed as sister group to *Haloarchaea*, members of the DPANN*-Archaea*, or sister group to *Methanocellales*. Screening a set of universal proteins, we identified four insertions located at critical locations in three ribosomal proteins and one RNA polymerase subunit that support the branching of *Nanohaloarchaea* as sister group to *Aenigmarchaea* within DPANN cluster II (*sensu* Dombrowski *et al*., 2020). Insertion analyses and phylogeny of the monomeric primase specific to DPANN*-Archaea* confirm the existence of a robust clade grouping *Undinarchaea*, *Naiadarchaea* and DPANN cluster II, that we propose to call *Nanostetteria*. Our insertion analysis also supports including *Altiarchaea* within DPANN-*Archaea* and suggest a new clades that has not been recovered in phylogenetic analyses, one grouping DPANN-*Archaea* with *Stygia* (*Hadarchaea* and relative) and an even large one grouping these lineages with *Acherontia* (*Thermococci* and relatives). The insertion defining this larger clade, present in the ribosomal protein uS7, is also present at the same position in *Thaumarchaea*, *Korarchaea* and a subgroup of *Asgardarchaea*. Whereas the insertion in *Thaumarchaea* is certainly due to an independent event, we discuss alternative hypotheses that can explain those present in *Korarchaea* and *Asgardarchaea.* Finally, we noticed several cases of MAGs misannotations, indicating that insertion analysis can be useful to identify protein with misleading affiliations. The existence of insertions in otherwise highly conserved universal proteins involved in translation or transcription could partly explain the high rate of protein evolution in some archaeal lineage, especially in DPANN*-Archaea*.

## Introduction

*Nanohaloarchaea* are tiny cellular organisms with small genomes that require high-salt concentrations for growth (Narasingarao et al., 2012, Hamm et al., 2019). They share their salty habitat with extremely halophilic archaea of the phylum *Euryarchaeota* (*Haloarchaea*) (Ghai et al., 2011, Narasingarao et al., 2012, Crits-Christoph et al., 2016, Hamm et al., 2019, La Cono et al., 2020, 2023, Najari et al., 2021, Baker et al., 2024). Despite the small sizes of their genomes, it was suspected for some time that *Nanohaloarchaea* could be free living organisms (Narasingarao et al., 2012). However, it was later shown that one of them, *Candidatus* Nanohaloarchaeum antarticus, can be only cultivated as an ectosymbiont of the haloarchaeum *Halorubrum lacusprofundi* (Hamm et al., 2019). Remarkably, *Candidatus* Nanohaloarchaeum antarticus fully internalizes within the *H. lacusprofundi* cytoplasm and subsequently trigger lysis of its host (Hamm et al., 2023). Similar to *Haloarchaea*, *Nanohaloarchaea* balance the extracellular high salt (sodium) concentration of their environment with high intracellular potassium concentrations (Narasingarao et al., 2012). Both *Haloarchaea* and *Nanohaloarchaea* are characterized by an acidic-rich proteome, allowing their proteins to remain soluble in their salt-rich cytoplasm. However, whereas *Haloarchaea* preferentially use aspartic acid to cover their protein surface with acidic charges, *Nanohaloarchaea* prefer glutamic acid (Narasingarao et al., 2012). *Nanohaloarchaea* were first positioned as sister group to *Haloarchaea* in an euryarchaeal phylogeny based on 16S rRNA (Narasingarao et al., 2012) and in an unrooted archaeal phylogeny based on conserved archaeal proteins (Petitjean et al., 2014) suggesting that adaptation to high salt concentration occurred only once in *Archaea* (Figure 1A). In contrast, *Nanohaloarchaea* were grouped with other nanosized archaea in phylogenies based on the concatenation of universal marker proteins, suggesting independent adaptation to high intracellular salt concentrations (Rincke et al., 2013, Williams et al., 2017, Castelle et al 2018) (Figure 1B). *Nanohaloarchaea* were elevated to the phylum rank (*Nanohaloarchaeota*) and included with other nanosized archaea in the proposed super-phylum DPANN (*Diapherotrites, Parvarchaeota, Aenigmarchaeota, Nanoarchaeota, Nanohaloarchaeota*) (Rincke et al., 2013). The grouping of *Nanohaloarchaea* with *Nanoarchaea* and *Parvarchaea* was also supported by a phylogeny based on the concatenation of 14 DNA replication proteins and by the replacement in these three lineages of the classical two subunits archaeal replicative DNA primase (PriS and PriL) by a smaller and divergent monomeric DNA primase, corresponding to a fusion of these two subunits (Raymann et al., 2014). Nowaday, it was also proposed that *Nanohaloarchaea* are neither DPANN nor sister group to *Haloarchaea* but are sister group to *Methanocellales*, that belong to *Methanomicrobia* (methanogens Class II) (Aouad et al., 2019) (Figure 1C). This result was based on a phylogenetic analysis of 68 core genes shared by *Nanohaloarchaea*, *Methanotecta* and *Diaphorarchaea* (these last two clades corresponding to group II *Euryarchaea, sensu* Forterre et al., 2014). The authors observed that *Nanohaloarchaea* branched in both ML and Bayesian trees at a position compatible with their grouping with other DPANN. However, when they used two independent strategies to reduce possible LBA artifacts (dayhoff4/6 recoding and the Slow-Fast method) *Nanohaloarchaea* became sister groups of *Methanocellales* (Figure 1C).

**Figure 1:**
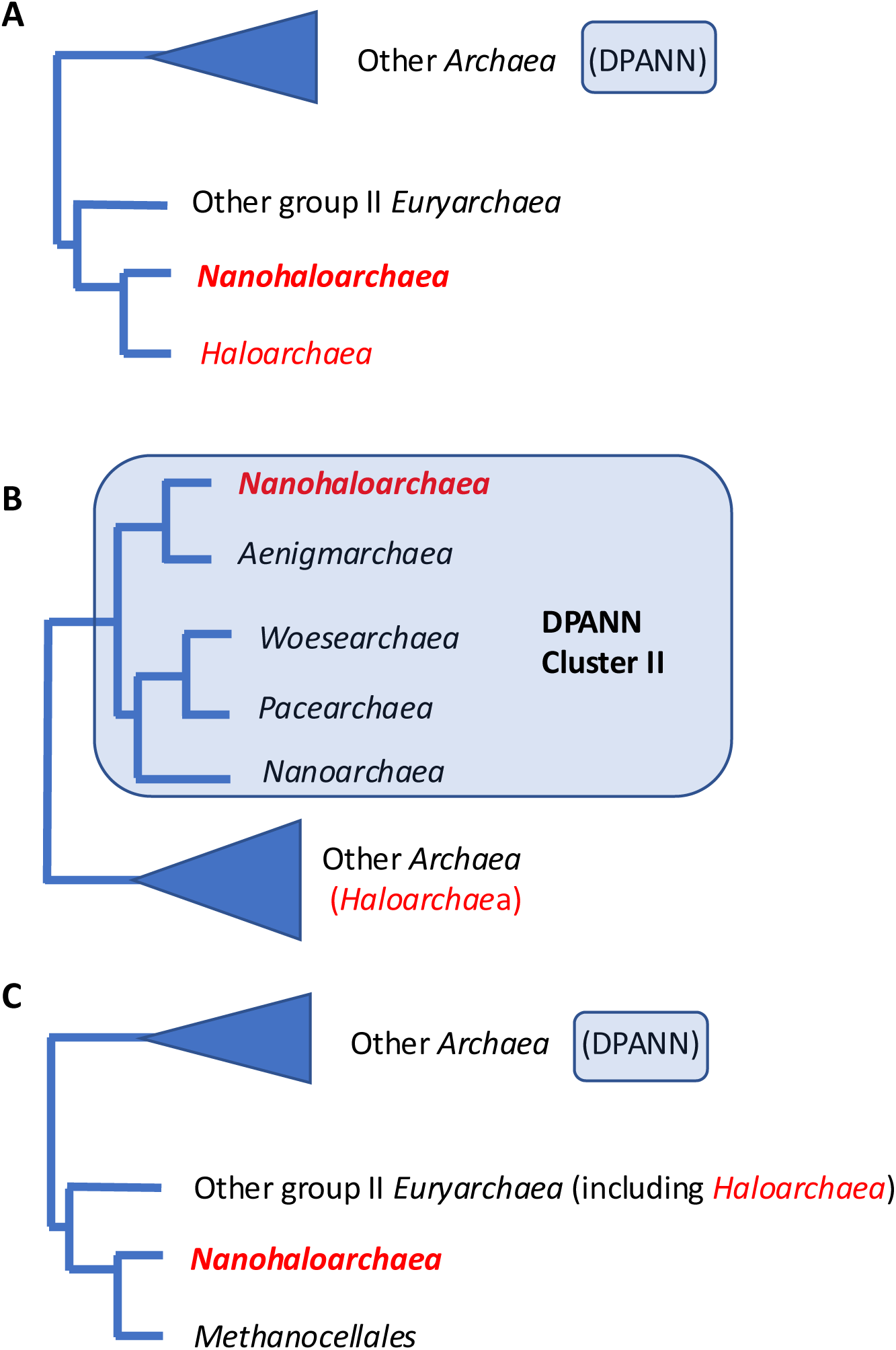
Different proposals made for the position of *Nanohaloarchaea* in the Archaeal tree. A, *Nanohaloarchaea* and *Haloarchaea* are sister group. Adaptation to high salt concentration was common to these two groups; B, *Nanohaloarchaea* belong to DPANN cluster II; C, *Nanohaloarchaea* are sister group to *Methanocellales*. In B and C, adaptation to high salt concentration occurred twice independently during the history of *Archaea*.

The proposed DPANN superphylum was expended with the identification of new lineages of nanosized archaea that also acquired the phylum status: *Woesearchaeota, Pacearchaeota, Micrarchaeota, Huberarchaeota* and *Undinarchaeota* (Castelle al., 2015, Schwank et al., 2018, Dombrowski et al., 2020). In a new nomenclature and taxonomy of *Archaea* recently implemented by the International Committee for the Systematic of Prokaryotes (ICSP), *Nanohaloarchaea* have conserved their phylum rank, whereas *Woesearchaeota, Pacearchaeota,* and *Parvarchaeota* have been included in the phylum *Nanoarchaeota*. More recently, all DPANN, including *Nanohaloarchaeota*, have been elevated to the kingdom status with the name, *Nanobdellati* (Göker and Oren, 2024), based on the characterization of a novel DPAN*N-Archaea*, *Nanobdella aerobiophila* as type species (Kato et al., 2022).

Considering that the taxonomic status and phylogenetic positions of many archaeal groups as well as the definition of phylum and superphylum are still debated (Panda *et al*., 2022, Lloyd and Tahon, 2022, Sutcliffe *et al*., 2021), here we will use the terms lineages with the suffix aea for the different groups under study, except for some well-defined classes, orders, or genera. We will also use the first proposed names for well-defined clades such as *Diapherotrites* (Rinke et al., 2013), *Stygia* (the clade grouping *Hadesarchaea* and *Persephonarchaea sensu* Adam et al., 2017) and *Acherontia* (the clade grouping *Thermococcales* with *Methanofastidiosa* and *Theionarchaea sensu* Adam et al., 2017). In the ICSP nomenclature, *Diapherotrites* correspond to the proposed phylum *Iainarchaeota*, *Hadesarchaea* correspond to the phylum *Hadarchaeota*, whereas the clades corresponding to *Acherontia* and *Stygia* have not been named (Rinke et al., 2021).

The grouping of *Nanohaloarchaea* with DPANN and the monophyly of DPANN were recently recovered in several phylogenies based on the concatenation of proteins conserved between DPANN and other *Archaea* (Schwank et al. 2019, Dombrowski et al., 2010, Aoud et al., 2022, Baker et al., 2024, 2025, Wen-Cong Huang, 2025). In the most recent analysis, the authors concluded that DPANN originated from free-living euryarchaeal ancestors (Baker et al., 2025).

However, the position of DPANN and *Nanohaloarchaea* in phylogenetic trees could be affected by the long branch attraction (LBA) artefact (Aouad et al., 2019, Feng et al., 2021, Rangel *et al*., 2021). DPANN indeed exhibit several characteristics of fast evolving parasites such as reduced genomes that correlated with the absence of otherwise essential metabolic pathways (for review see Dombrowski et al., 2019). Strikingly, DPANN were either recovered at the base of the archaeal tree in most archaeal phylogenies using *Bacteria* as the outgroup (Hug et al., 2016, Williams et al., 2017, Liu et al., 2021, Martinez-Guiterrez and Aylward, 2021, Moody et al. 2022), suggesting their attraction by the long branch of *Bacteria*.

Remarkably, DPANN are still divided into eleven phyla in the ICSP nomenclature, whereas *Asgardarchaea* and TACK-*Archaea* are each reduced to one phylum, *Thermoproteota* This suggests that the bias introduced by the fast-evolving character of DPANN proteins was still present in the phylogenetic analysis used by Hugenholtz and collegues to calculate the relative evolutionary divergences (RED) at the base of their ranking method and of the recent ICSP nomenclature (Rinke et al., 2021; Göker and Oren, 2024).).

The grouping of *Nanohaloarchaea* among DPANN has been especially controversial since, in addition to their fast-evolving character, they exhibit the atypical amino acid composition of halophilic proteins and could have experienced extensive lateral gene transfer (LGT) with their haloarchaeal hosts (Rangel et al., 2021, Baker et al., 2025). The problem was specifically addressed by Gogarten and colleagues who analyzed four different protein datasets, the ATP synthase catalytic and non-catalytic subunits, a concatenation of 44 ribosomal proteins and two core protein datasets containing 12 and 146 proteins, respectively (Feng et *al*., 2021). They obtained the grouping of *Nanohaloarchaea* with DPANN in their ribosomal and core protein phylogenies using species dataset containing several DPANN species. However, they obtained the sisterhood of *Nanohaloarchaea* and *Haloarchaea* with their ATP synthase phylogeny, but also with their ribosomal and core protein phylogenies when they removed other DPANN archaea from their species dataset (Feng *et al.,* 2021). Finally, Fournier and colleagues identified in Archaea different “clusters” of genes that produced coherent phylogenies within each cluster but different ones between them. When they analyzed the position of *Nanohaloarchaea* in the absence of other DPANN, two of these clusters grouped again *Nanohaloarchaea* and *Haloarchaea* together (Rangel et al., 2021).

Here, we have revisited the problem of the *Nanohaloarchaea* position by analysing insertions in the alignment of universal proteins that could be used as synapomorphies to discus this position. In some cases, the analysis of insertions can be useful to confirm or not the validity of clades identified by classical phylogenetic analysis. For instance, classical phylogenetic analysis, of the EF1-alpha elongation factor of *Microsporidia* provide trees in which these unicellular eucaryotes without mitochondria were located at the base of the eucaryotic tree, supporting the archezoa hypothesis (Kamaishi et al., 1996). However, it was also noticed that the sequence of the EF1-alpha elongation factor of microsporidia share a long 15 amino acids insertion with fungi (Kamaishi et al., 1996), and it was clearly demonstrated later on that microsporidia are a subgroup of fungi (James et al., 2006).

We report here the identification of four insertions that support the grouping of *Nanohaloarchaea* (including the recently described *Asbonarchaeaceae*) with a subgroup of DPANN (DPANN cluster II, *sensu* Dombrowski *et al*., 2020) as sister group to *Aenigmarchaea*. Our analysis confirms the existence of a clade grouping all DPANN encoding the monomeric primase that we propose to call *Nanostetteria*. We also confirm the close relationships between *Altiarchaea* and DPANN. Interestingly, our indel analysis suggests the existence of a previously unrecognized clade grouping DPANN, *Altiarchaea*, *Stygia* (*Hadesarchaea* and *Persephonarchaea*), and *Acherontia* (*Thermococcales, Methanofastidiosa* and *Theionoarchaea*) thereafter called the DASA clade. Notably, similar insertions to one of those analyzed here are present in a subgroup of Asgard, suggesting to divide Asgard in two major clusters and raising the possibility of an unexpected evolutionary connection between Asgard and this DASA clade. We conclude that insertion analysis can be useful to unravel the position of archaeal species with unstable position in classical phylogentic analysis. The insertions analyzed here are present in conserved and critical regions of ribosomal proteins and RNA polymerase sequences, suggesting that alteration of transcription and translation could partly explain the fast-evolving character of DPANN proteins, and raising the possibility that other archaeal lineages, including Asgard, are also fast evolving organisms. Finally, we detected during our analyses several cases of MAG misannotations, indicating that insertion analysis can be also useful to identify problems in metagenome reconstruction.

## Results

We have previously analyzed the phylogenies of the 36 universal proteins used by Spang and colleagues (Spang et al., 2015, 2018) to determine the position of *Asgardarchaea* in the tree of life (Da Cunha *et al*. 2017, 2018). In the course of this work, we noticed that the sequences of several archaeal lineages were enriched in insertions or deletions (indels) compared to other lineages (Table S1). This was especially the case for sequence of the DPANN *Nanoarchaeum equitans* (see an example in Figure S38 of Da Cunha *et al*. 2017) in agreement with the idea that these episymbionts with reduced genome are fast-evolving organism (Brochier *et al*., 2005). We had previously observed this phenomenon in studying the sequence of the *Methanopyrus kandleri* RNA polymerase large subunits, since we observed that these sequences have accumulated 27 insertions *versus* 1 to 8 for RNA polymerases of the other archaeal lineages known at that time (see Table 1 in Brochier *et al*., 2004). We analyzed in more details some of the specific insertions present in *N. equitans* and and we noticed that several of these indels were present in *Nanohaloarchaea*. We thus reasoned that screening systematically for insertion in universal proteins of *Nanohaloarchaea* could be a wise strategy to check the position of these organisms in the archaeal tree. After adding several sequences of DPANN-*Archaea* (including two *Nanohaloarchaea*), *Altiarchaea* and *Asgardarchaea* in our previous alignments of the 36 proteins dataset (Da Cunha et al., 2017) and the alignment of the ribosomal protein uS5 (see Materials and Methods), we identified four insertions conserved in all *Nanohaloarchaea* that have taken place in regions otherwise very well conserved in the three domains or at least between *Archaea* and *Eukarya*. These insertions were present in all sequences of Nanohaloarchaea (around 50) present in the non redundant (nr) protein sequence database of the NCBI in march 2026. Three of these insertions are present in the ribosomal proteins uS5, uS7 and uL16 and a fourth one in the B” subunit of the DNA-dependent RNA polymerase. All insertions detected here are located between regions of highly conserved amino-acid signatures that can be used as anchors to validate the alignments. They are also all located in regions important for the function of the proteins in translation or transcription (see below).

### Large insertions in the ribosomal proteins uS5 and uS7 indicate that *Nanohaloarchaea* are members of a clade grouping seven lineages of DPANN-*Archaea* cluster II

In uS5 (previously named S2e in eucaryotes), an insertion of 15 amino acids is present in all *Nanohaloarchaea* (four examples in the red frame of Figure 2) in a region where no insertion is present in the three domains of life, including in the fast evolving archaeal species such as *Methanopyrales* and *Korarchaea* (blue arrows). A similar insertion is only present in several other DPANN lineages (see below). The superimposition of an AlphaFold (Jumper et al., 2021) model of uS5 from *Candidatus* nanohaloarchaeota archaeon to uS5 in the *P. abyssi* 30S structure shows that this insertion affects a β-hairpin that protrudes into the mRNA entry channel (Figure 3). uS5 and its β-hairpin region are known to be involved in start codon selection (Dong and Hinnebusch, 2022) and translational fidelity (Kamath et al., 2017). Remarkably, insertions of the same size (with six strictly conserved amino acids at the same position, G…GK.GG..R) are present in all *Aenigmarchaea, Pacearchaea, Parvarchaea, Woesearchaea,* and *Huberarchaea* (five examples in red frame Figure 2). The insertion present in *Nanohaloarchaea* is especially similar to those of *Aenigmarchaea,* with 8 strictly conserved amino acids (GG.PGKGGG..RT). A slightly smaller insertion of 13 amino acids with a rather similar sequence (containing the dinucleotide GK) is present at the same position in the uS5 protein of *N. equitans* and other well characterized genera of *Nanoarchaea* such as *Nanopusillus*, *Nanobdella aerobiophila*, or else *Candidatus* Nanobsidianus stetteri (2 example in Figure 2). All these lineages indeed form a clade in most classical phylogenetic analyses and have been grouped under the name DPANN cluster II (black frame in Figure 2) by Spang and colleagues (Dombrowski et al., 2020). The 13-amino-acid insertion of Nanoarchaea could be useful to distinguish *bona fide* relatives of *N. equitans* from other DPANN that are annotated as “Nanoarchaeota archaeon” in the NCBI database Two very similar smaller insertion of three and four amino-acids are also present at the same position in the uS5 protein of *Undinarchaea* and *Naiadarchaea* (red arrow) confirming that these two lineages forms a distinct clade among DPANN. One of these insertions includes the GKG motive that can be aligned by hand with the GKG of *Aenigmarchaea* and *Nanohaloarchaea* (not shown).

**Figure 2:**
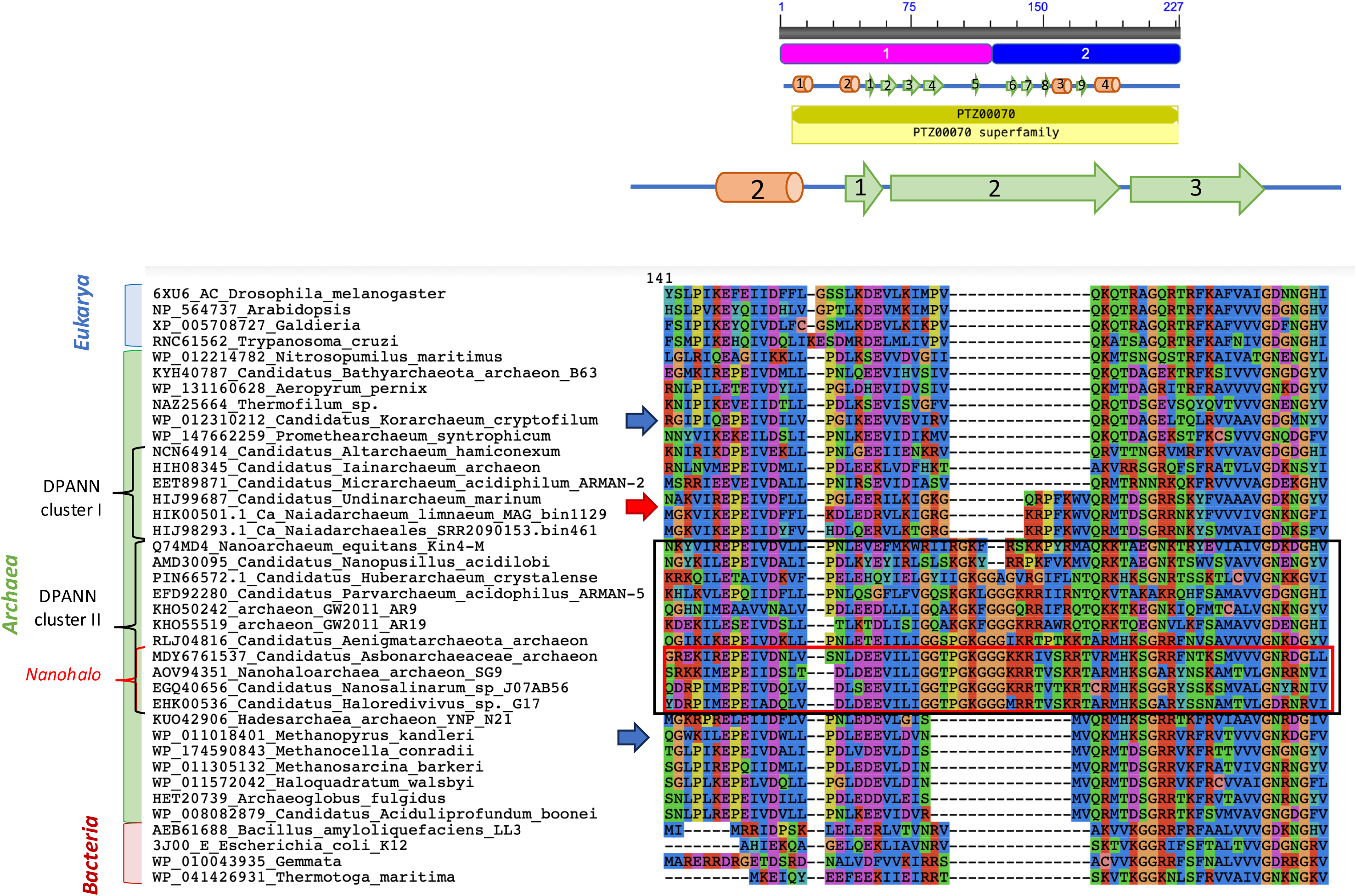
The uS5 insertion: Alignment of selected sequences of the uS5 insertion. Sequences of *Nanohaloarchaea* are in the red frame and those of DPANN cluster II in the black frame. Sequences of the fast evolving species *Korarchaeaum cryptophilum* and *Methanopyrus kandleri* are indicated by blue arrows. The sequence of an Undinarchaeon is indicated by a red arrow.

**Figure 3:**
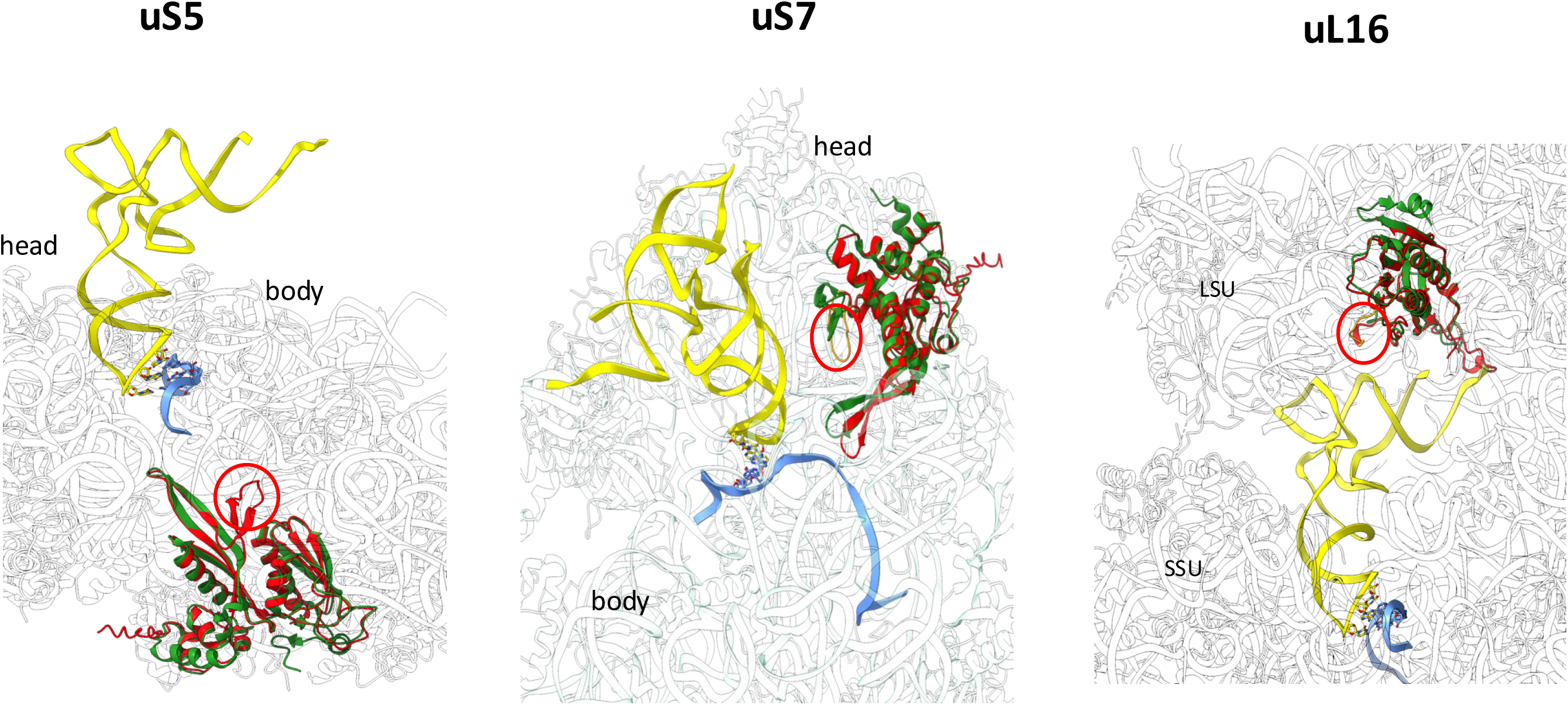
**Localization of the insertions detected in ribosomal proteins. uS5**: uS5 in a *P. abyssi* 30S initiation complex (PDB ID Code 7ZAH, Kazan et al 2022, PMID: 35694843). The P site initiator tRNA is in yellow, the mRNA is in blue, uS5 is in green. In red, the AlphaFold model of uS5 from a *Candidatus* Nanohaloarchaeota archaeon superimposed to *P. abyssi* uS5. The insertion described in Figure 2 is circled in red. **uS7**: uS7 in a *P. abyssi* 30S initiation complex. The initiator tRNA bound to the P site is in yellow, the mRNA is in blue, uS7 is in green. In red, the AlphaFold model of uS7 from *Candidatus* Nanohalobium constans superimposed to *P. abyssi* uS7. The insertion described in the upper panel is in orange and circled. **uL16**: uL16 in *T. kodakarensis* 70S ribosome (PDB ID Code 6TH6, Sas-Chen 2022, PMID: 32555463) with mRNA (blue) and P site tRNA (yellow) from P. abyssi (PDB ID Code, 7ZAH (Kazan et al 2022, PMID: 35694843). uL16 from *T. kodakarensis* is in green. In red, the alpha-fold prediction of uL16 from *Candidatus* Nanohalobia archaeon BNXNv superimposed to *P. abyssi* uS7. The insertion described in the upper panel is in orange and circled. Large (LSU) and small (SSU) ribosomal subunits are indicated.

The absence of insertion at this position in other archaea classified as DPANN, i.e. *Micrarchaea*, *Diapherotrites* (*Ianarchaea*), *Altiarchaea*, strongly suggests that this insertion corresponds to a synapomorphy testifying for a clade grouping all members of DPANN cluster II with *Undinarchaea* and *Naiadarchaea* (Dombrowski et al. 2020) (black frame in Figure 2).

In uS7 (previously named S5e in *Eukarya*), an insertion of 11 amino acids is present in all *Nanohaloarchaea* (5 examples in red frame in Figure 4A) in a region which is conserved in size in the three domains of life, except for several groups of *Archaea*, including DPANN, *Altiarchaea*, *Stygia, Acherontia, Korarchaea* and *Thaumarchaea*. Based on the superposition of an AlphaFold model of *C.* nanohalobium constans uS7 with the *P. abyssi* 30S structure (Figure 3), this insertion is located close to an rRNA region universally involved in the selective binding of the initiator tRNA (Coureux et al., 2020).

**Figure 4:**
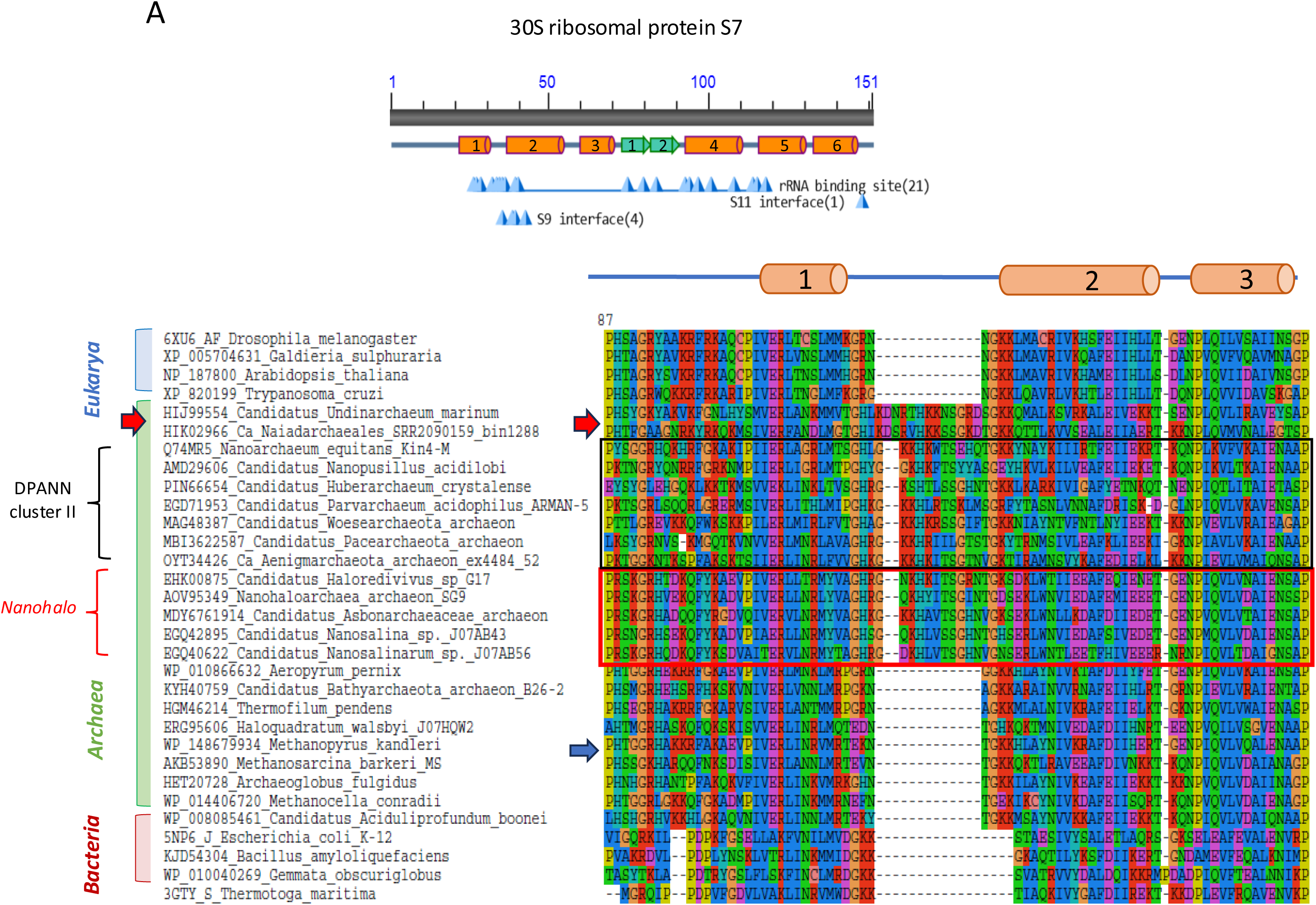
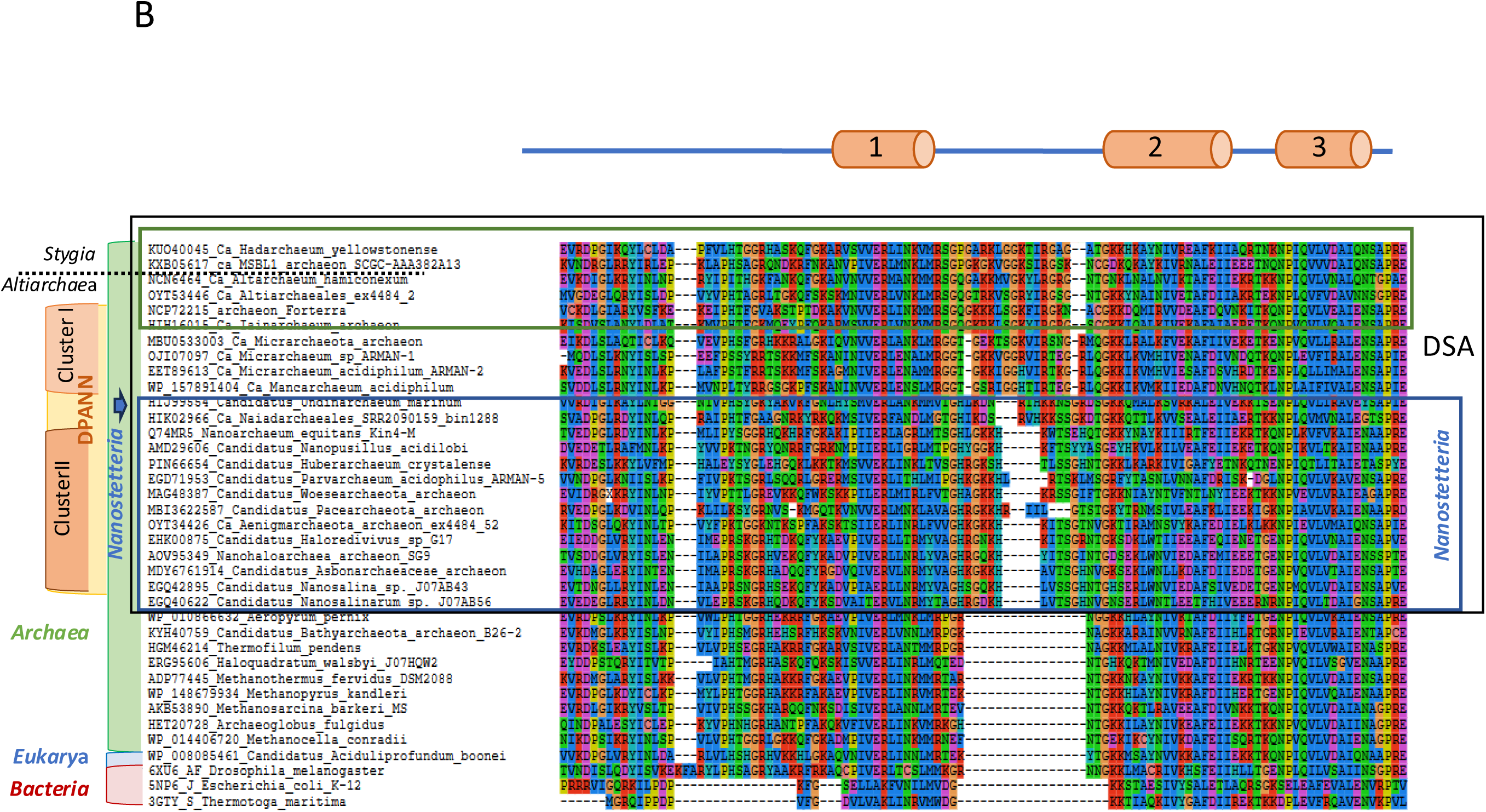
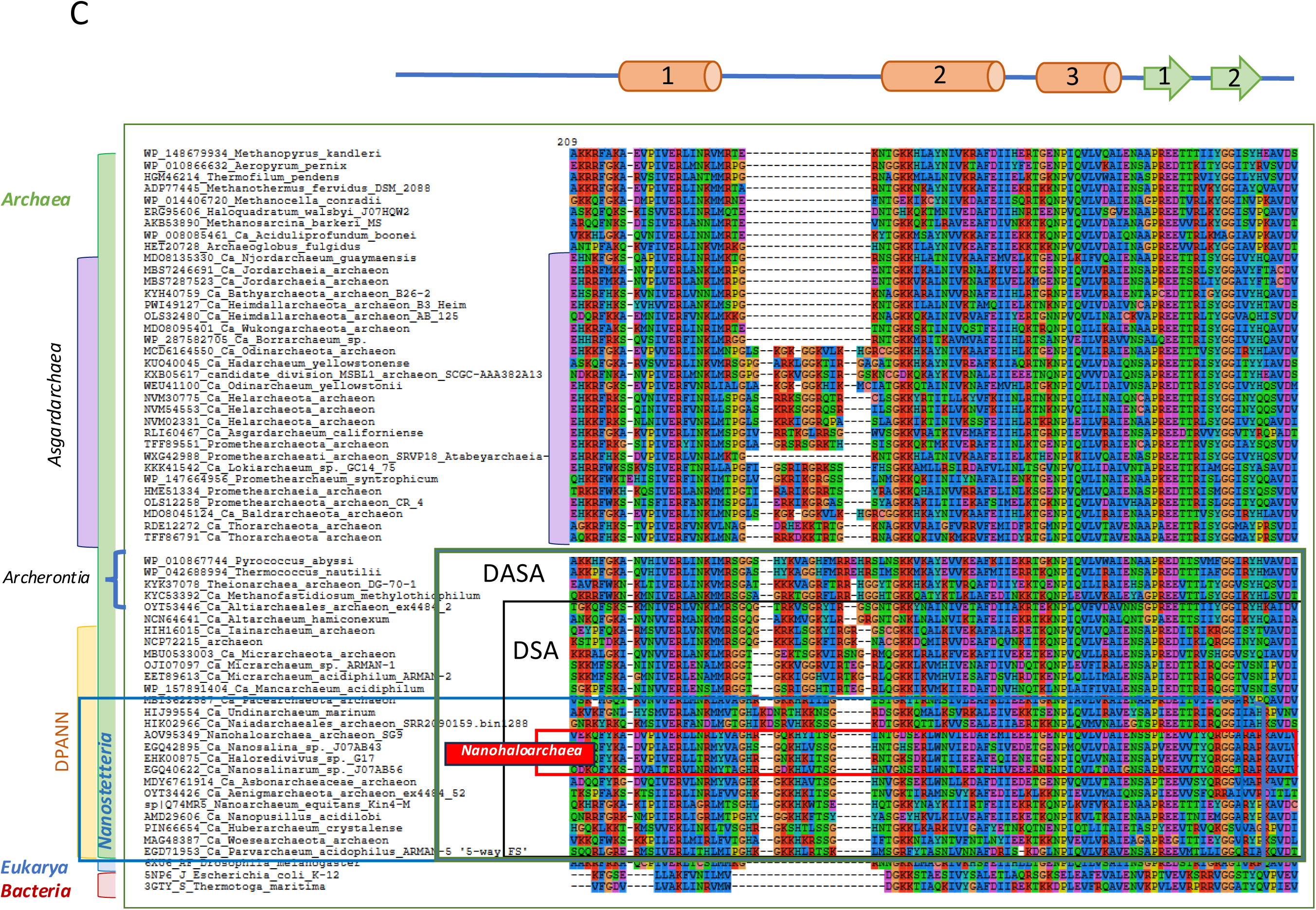
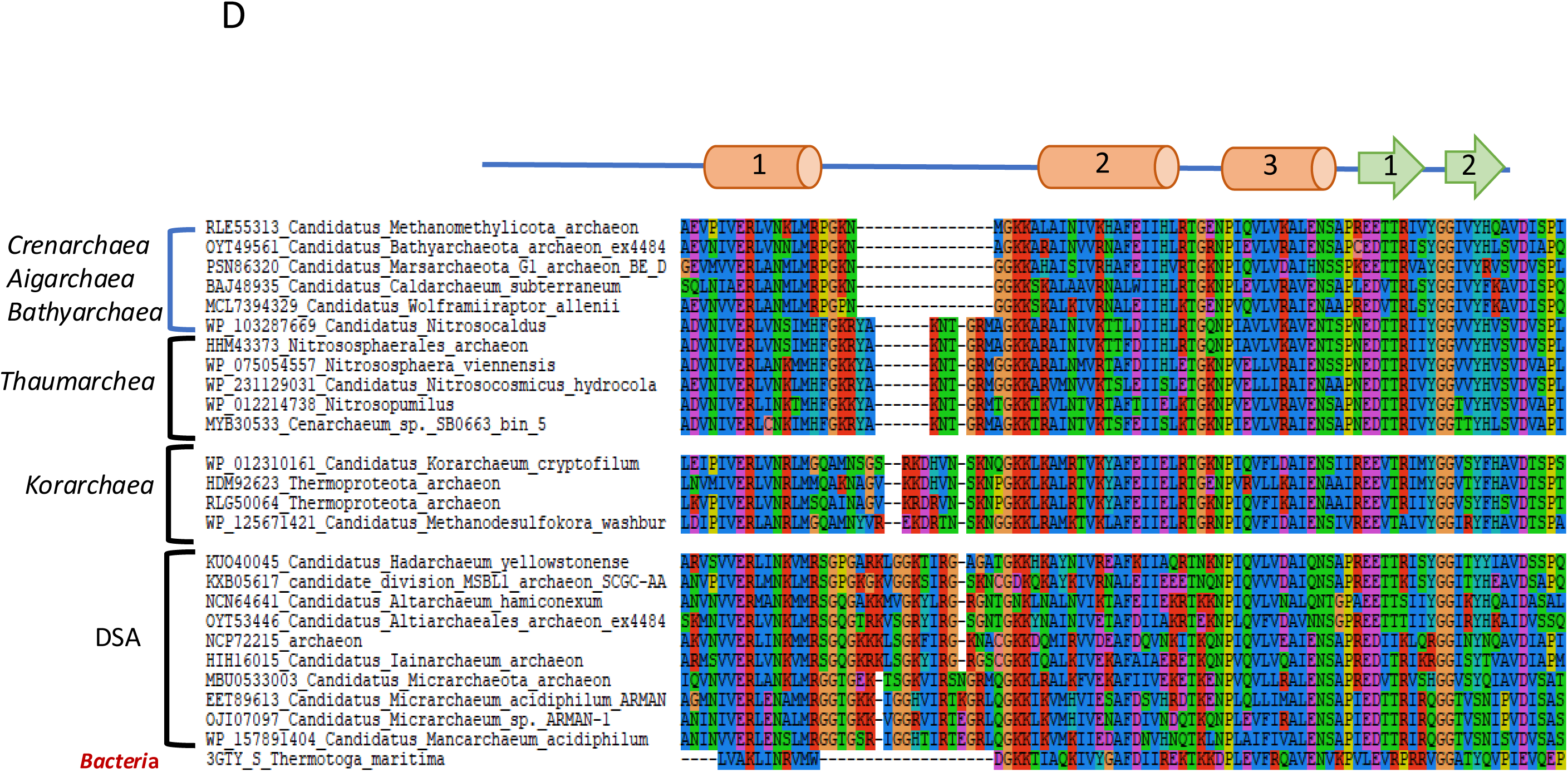
**The uS7 insertion**: 4A: Alignment of selected sequences of the uS7 insertion in *Nanohaloarchaea*, DPANN cluster II, *Undinarchaea* and *Naiadarchaea* corresponding to the proposed clade *Nanostetteria* with representatives of selected groups that do not contain this insertion. Sequences of *Nanohaloarchaea* are in the red frame and those of DPANN cluster II in the black frame. Blue arrows indicate the known fast evolving species *M. kandleri*. 4B: Alignment of selected sequences of the uS7 insertion of *Nanostetteria* (blue frame) with a selection of insertions present in DPANN cluster 1, *Altiarchaea* and *Stygia*. Blue arrow indicates the position of *Uninarchaea* and *Naiadarchaea* that have been included in *Nanostetteria*. Note that the perfect alignment of DPANN cluster II sequences and their alignment with *Uninarchaea* and *Naiadarchaea* in Figure 4A has been slightly modified by the addition of new sequences. Figure 4C: Alignment of a selection of the insertions present in *Nanostetteria*, DPANN cluster 1, *Altiarchaea* and *Stygia* with a selection of insertions present in *Acherontia* and *Asgardarchaea*. Figure 4D: tentative alignments of the uS7 insertions present in DPANN and *Stygia* with those present in *Korarchaea* and *Thaumarchaea*.

Strikingly, 11 amino-acids insertions very similar to those of *Nanohaloarchaea* with the strictly conserved signature signature GxKH are only present in members of DPANN cluster II (black frame in Figure 4A). A 13 amino-acid insertion was also found at this position in *Undinarchaea* and *Naiadarchaea* (red arrows in Figure 3). This insertion could be aligned with those of DPANN cluster II with conservation of the HK and SG motives motives, indicating that they are most likely homologues. Both the uS5 and uS7 insertions thus indicate a close relationships between DPANN cluster II, *Undinarchaea* and *Naiadarchaea*, these two lineages being probably sister group, in agreement with recent phylogenetic analyses (Dombrowki et al. 2019).

Notably, the DPANN lineages sharing the uS5 and uS7 insertions, including *Undinarchaea* and *Naiadarchaea*, are the only lineages that share the unique monomeric primase previously detected in DPANN archaea (Raymann et al., 2014, Dombrowski et al., 2020). This strongly suggests that this subgroup of DPANN archaea form a monophyletic group including DPANN cluster II, *Undinarchaea* and *Naiadarchaea*, as previously observed in the phylogenies published by Spang and colleagues (Dombrowski et al., 2020). It makes sense to give a name to this robust, well-defined clade. We suggest calling here members of this clade *Nanostetteria* considering that *N. equitans,* episymbiont of the crenarchaeon *Ignicoccus equitans,* has been the first identified and isolated in the laboratory of Karl Stetter (Huber et al., 2002).

### Larger similar uS7 insertions are shared by Nanostetteria, DPANN cluster I, Stygia, and Acherontia

Large insertions of 14 amino acids with significant sequence similarities to the 11-amino acid insertion of *Nanostetteria* are present in the same region of the uS7 protein in DPANN cluster I (*Diapherotrites/Iainarchaea, Micrarchaea*), *Altiarchaea* and *Stygia* (MSBL *Persephona, Hadesarchaea /Hadarchaea*) (black frame in igures 4B). All these insertions were rather well aligned with those of the DPANN cluster II by the program used (see Materials and Methods). Overall, the examination of the alignment in Figure 4B and of various possible alternative hand-made alignments (not shown) suggest that all these insertions are homologous. This supports the existence of a large clade grouping *Nanostetteria,* DPANN cluster I and *Altiarchaeales* with *Stygia* (*Hadarchaeota*). We will call thereafter this putative clade, the DSA clade for DPANN, *Stygia, Altiarchaea*.

These 14 amino-acid insertions are especially similar in *Diapherotrites/ Ianarchaea*, *Altiarchaea*, *Stygia,* and in the *Micrarchaea* labelled as *Forterra,* with the conserved 5-amino acid motif GxxxxxGxxIRG, whereas the final IRG motif is transformed into IRxxG in other *Micrarchaea*. This confirms the close relationships between DPANN and *Altiarchaea* (Kellner et al., 2018, Dombrowski et al., 2020, Castelle et al., 2021, Rinke et al., 2021, Aouad et al., 2022, Baker et al., 2025) and suggests to divide the DSA clade into three subclades, one corresponding to *Nanostetteria* (blue frame), another grouping DPANN cluster I, *Stygia*, *Altiarchaea* and *Micrarchaea* related to *Forterra* (green frame I Figure 4B) and a third one grouping other *Micrarchaea*. If this interpretation:n is correct, it means that *Micrarchaea* are paraphyletic and thus do not correspond to a valid clade.

The 14-amino acid insertions present in members of the DSA clade (black frame in Figure 4C) were aligned with a 15-16 amino acids insertion present at the same position in Acherontia (*Thermococcales* and relatives, 4 examples in Figures 4C). All these insertions were aligned around a conserved glycine. These insertions are well conserved between *Altiarchaea*, *Stygia* and *Acherontia* with the conserved motif KxxGxxxR, suggesting that all these insertions are homologous. A specific evolutionary connection has been previously suggested between *Nanoarchaea* and *Thermococci* based on phylogenetic analyses (Brochier et al., 2004). It is tempting to suggest that the insertions detected here in uS7 testifies for a large superclade grouping all DPANN, *Altiarchaea, Stygia* and *Acherontia* (therafere called here the DASA clade for DPANN, *Altiarchaea, Stygia* and *Acherontia*) (green frame in Figure 4C). Note that addition of sequences in Figure 4B and C has slightly modified the perfect alignment of the insertions present in the *Nanostetteria* uS7 proteins, indicating that better alignments of insertions are obtained inside a specific clade when the aligned sequences are restricted to this clade and a close outgroup missing the insertions.

### Larger uS7 insertions are also present in a subgroup of *Asgardarchaea*

One of us (PF) has recently described large insertions (10-13 amino-acids) in the uS7 proteins of a subgroup of Asgardarchaea (*Lokiarchaea, Helarchaea, Thorarchaea, Baldrarchaea, Hermodarchaea* and *Odinarchaea*) (Figure S2 in Forterre, 2025). The lineages of *Asgardarchaea* containing these insertions form a monophyletic group in the concensus phylogenies of *Asgardarchaea* that we recently proposed (Da Cunha et al., 2022, Forterre, 2025). Notably, these insertions are located at the same position than the insertions typical of the DASA clade (Figure 4C). When we added several sequences of *Asgardarchaeota* to our alignments, the insertions present in both DASA and those *Asgardarchaea* were aligned by the program based on a conserved glycine present in the center of these insertions, except in some *Thorarchaea* (25 examples of Asgard in Figure 4C). The sequences of these insertions are well conserved within lineages of *Asgardarchaea* but specific for each of them, indicating that this region changes rather rapidly in this group. Considering their low sequence similarity with the DASA insertions, these insertions could have occurred independently in DASA and *Asgardarchaea*, despite their similar size and location. However one cannot exclude the possibility of an unexpected evolutionary connection between DASA and this subgroup of *Asgardarchaea*.

### Larger uS7 insertions are also present in *Thaumarchaea* and *Korarchaea*

Insertions of 8 and 12 amino acids are also present in *Thaumarchaea* and *Korarchaea* in uS7 proteins at the position of the insertions present in DASA members (Fig. 4D). They exhibit less obvious similarities with the DASA insertions, except for an enrichment in basic amino acids. The 8 amino acid insertion, which is conserved in all *Thaumarchaea,* is absent in *Bathyarchaea* and *Aigarchaea*. *Thaumarchaea* form with *Bathyarchaea* and *Aigarchaea* a robust clade (the BAT clade according to Gaïa et al., 2018) which supported in most phylogenetic analyses (Adam et al., 2017, Da Cunha et al., 2017, Jay et al., 2018, Rincke et al., 2021, Aouad et al., 2022). This strongly suggests that the 8 amino-acid insertion has been introduced in the lineage leading to *Thaumarchaea* independently of those present in the DASA clade. The 12 amino-acid insertions present in *Korarchaea* have also few sequence similarities with those of DASA members, suggesting that the presence of insertions at similar position in DASA*-archaea*, *Thaumarchaea* and *Korarchaea* is more likely due to convergent evolution driven by physical constraints determining the nature of the amino acids present in these insertions. However, the sizes of these insertions being rather similar in *Korarchaea*, *Asgardarchaea* and DASA*-Achaea*, and the position of *Korarchaea* in the archaeal tree being still unresolved (Liu et al., 2022), open cannot exclude a more complex evolutionary scenario (see discussion).

In the examples shown in Figure 4D, it is interesting to note that two sequences of *Korarchaea* are annotated as *Thermoproteota* archaeon, which is not informative since this putative phylum in the ICSP nomenclature includes many other former phyla such as *Crenarchaea, Marsarchaea, Methanomethylicaea, Thaumarchaea*, *Aigiarchaea* or else *Bathyarchaea*. In that case, this specific insertion can help to identity *Korarchaea* from other “Thermoproteota”. Notably, the regions bordering the left side of the 12 amino acids insertion in *Korarchaea* were difficult to align with the corresponding regions in uS7 proteins from the three domains of life, indicating a higher variability of *Korarchaea* in that region. For instance, a universally conserved glycine is absent in *Korarchaea*.

### A uL16 insertion suggests including *Altiarchaea* within DPANN-*Archaea*

The protein uL16 (formerly called L10e in eukaryotes) of all *Nanohaloarchaea* contains a 6/7-amino acid insertion. This insertion is present in a region which is highly conserved in *Eucarya* and *Archaea*. We superposed an AlphaFold model of a *Candidatus* nanohalobia archaeon uL16 on the *T. kodakarensis* 70S structure (Sas-Chen et al., 2020). The insertion is located close to the acceptor stem of the P site tRNA (Figure 3). Similar insertions, most likely homologous, are present in other *Nanostetteria*, but also in all DPANN cluster I (eight example in Figure 5) and in all *Altiarchaea* (2 examples in Figure 5). This confirms the close relationships between *Altiarchaea* and DPANN, as previously observed in the case of the uS7 insertion and in several phylogentic analyses. The absence of this insertion in *Stygia* support the existence of a clade grouping *Nanostetteria*, DPANN cluster I and *Altiarchales*, suggesting to conside *Altiarchaea* as members of DPANN-*Archaea*. One of us (PF) previously detected a 6/9 amino acid insertion at the same position in a subgroup of Asgard tht includes *Lokiarchaea, Helarchaea* and *Odinarchaea* (Forterre, 2025). Since this insertion is absent in all other *Asgardarchaea,* but also in *Stygia* and *Acherontia*, it is likely that the insertions in DPANN-*Archaea* and in this subgroup *Asgardarchaea* were independent events.

**Figure 5:**
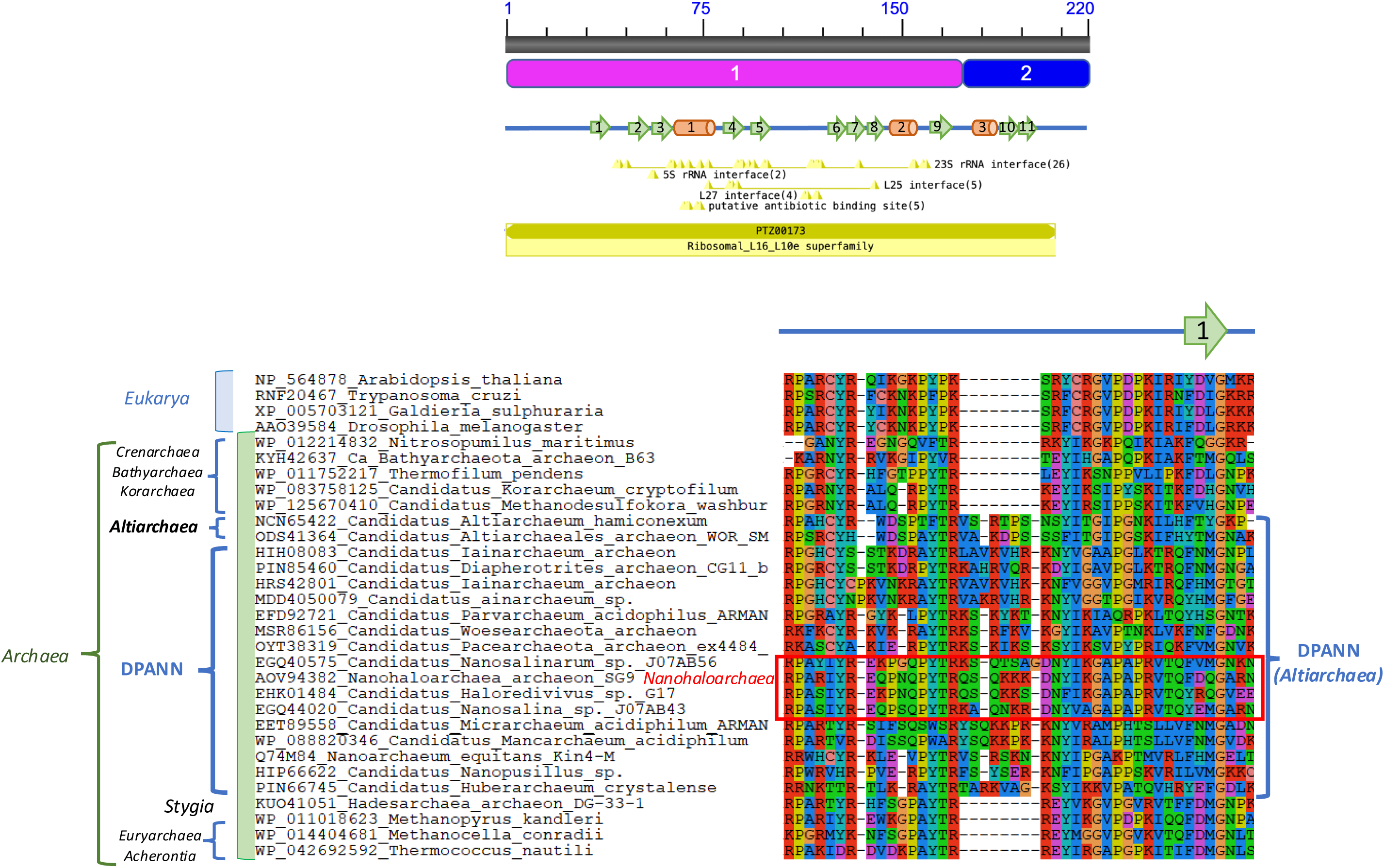
**The uL16 insertion**. Alignment of selected sequences of the uL16 insertion present in DPANN *Archaea* and *Altiarchaea* with selected sequences from other *Archaea* and *Eukarya*.

### An insertion in the RNA polymerase B” subunit supports the clade grouping Nanohaloarchaea *with* Aenigmarchaea

All *Nanohaloarchaea* share a 3amino acid insertion in the protrusion domain of the RNA polymerase B” subunit (red frame in Figure 6A) (the B subunit is split into B’ and B” subunits in all *Euryarchaea* and most DPANN). This insertion is located within an α helix that belongs to the protrusion domain of the B subunit (Bushnell et al., 2002, Cramer et al., 2016). This region is conserved in the three domains of life and should be important for the function of the RNA polymerase. The region containing this insertion is conserved in all other archaeal RNA polymerases, except for a similar insertion in *Aenigmarchaea*. The insertions in *Aenigmarchaea* is similar in size and sequence to those in *Nanohaloarchaea*, supporting again the sisterhood of these two lineages within DPANN cluster II.

**Figure 6:**
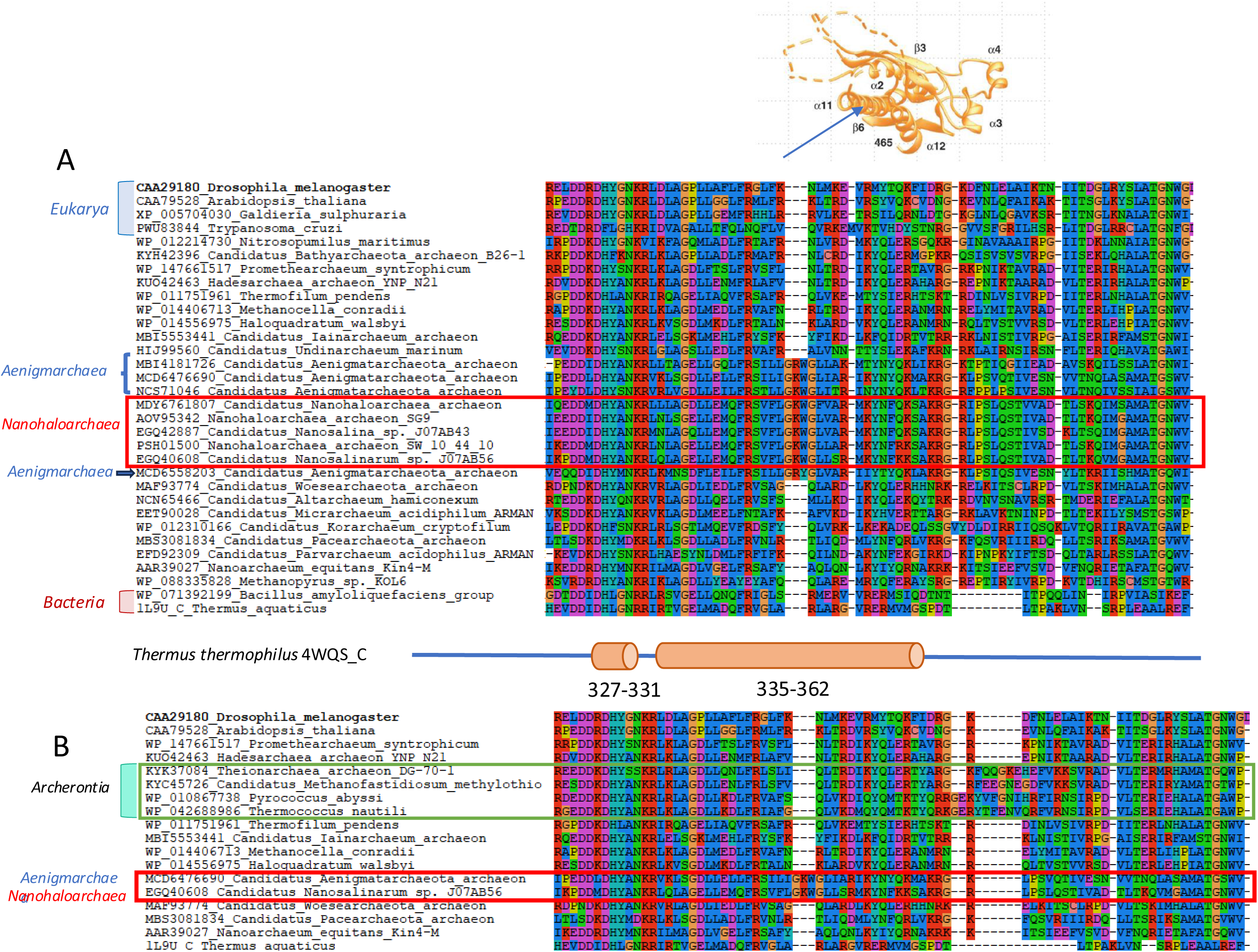
RNA polymerase B” insertion. A) Upper figure: structure of the RNA polymerase B region containing the specific insertion present in *Aenigmarchaea* and *Nanohaloarchaea*. The insertion occurred in the α11 helix (from Cramer et al., 2016). A: Alignment of selected sequences containing the RNA polymerase B” subunit insertion with selected sequences of corresponding regions of RNA polymerases B or B” sequences that do not contain this insertion. B: Alignment of selected sequences of RNA polymerase B subunit containing or not the insertion present in *Aenigmarchaea* and *Nanohaloarchaea* (framed in red) with selected sequences of *Acherontia* (framed in green). Sequences corresponding to insertions are overlined in yellow.

Interestingly, we noticed on the right side region of this insertion a 9 amino acids insertion in *Thermococcales* and a 7 amino acids insertion in *Theinoarchaea* and *Methanofastidiosa* that are located at the same position and could be synapomorphy supporting the clade *Acherontia* (green frame in Figure 6B) (Adam et al., 2017).

### Phylogeny of DNA primase

*Nanohaloachaea* and several other DPANN lineages lack the classical two-subunits eucaryotic-like archaeal DNA primase (PriS, PriL) and encode instead for a very divergent monomeric DNA primase (Raymann et al. 2014, Dombrowski et al., 2020). This DNA primase is formed by the fusion of PriS and PiL domains, ressembling DNA polymerase/Primase encoded by some archaeal plasmids (Gill et al., 2014). Notably, if *Nanohaloarchaea* do not belong to DPANN, the presence of this unique DNA primase in *Nanohaloarchaea* can be only explained by independent acquisitions of homologous atypical DNA primases (probably from mobile elements) and/or LGT between different DPANN lineages. However, this hypothesis is not trivial, since *Nanohaloarchaea* and other DPANN containing this primase did not share similar biotopes. It seems also unlikely that they share similar types of mobile elements if they are phylogenetically distant, since archaeal mobile elements usually co-evolve with their hosts in *Euryarchaea* (Forterre et al., 2014).

To explore the distribution and evolutionary history of the *Nanostetteria* DNA primase, we first recovered all sequences homologous to the DNA primase of *N. equitans* and we built a phylogeny of these DNA primases using Maximum likelihood inference with IQtree (see Materials and Methods). We only found homologous DNA primase in *Nanostetteria*, with few exceptions (including two *Bacteria*) that most likely correspond to contaminations or recent gene transfer. For instance, we detected the gene encoding the monomeric DNA primase in two MAGs annotated as *Candidatus* bacterial sequences whereas these two primases branched within *Woesearchaea* in phylogenetic analyses, or else, we detected monomeric DNA primase in MAGs annotated as *Micrarchaea* and *Altiarchaea*, whereas their sequences branch within *Aenigmarchaea* in our DNA primase phylogenies. We also noticed some case of MAGs annotated as *Nanoarchaeota* whose primases branch far from *bona fide N. equitans* and other *Nanoarchaea* whose genomes have been completely sequenced. Finally, we detected several clades of DNA primases that could correspond to yet non-identified *Nanostetteria* lineages. All these sequences were removed from our final phylogenetic analysis (see materials and methods) to get a simplified but accurate version of the DNA primase history (Figure 7 and S2).

**Figure 7:**
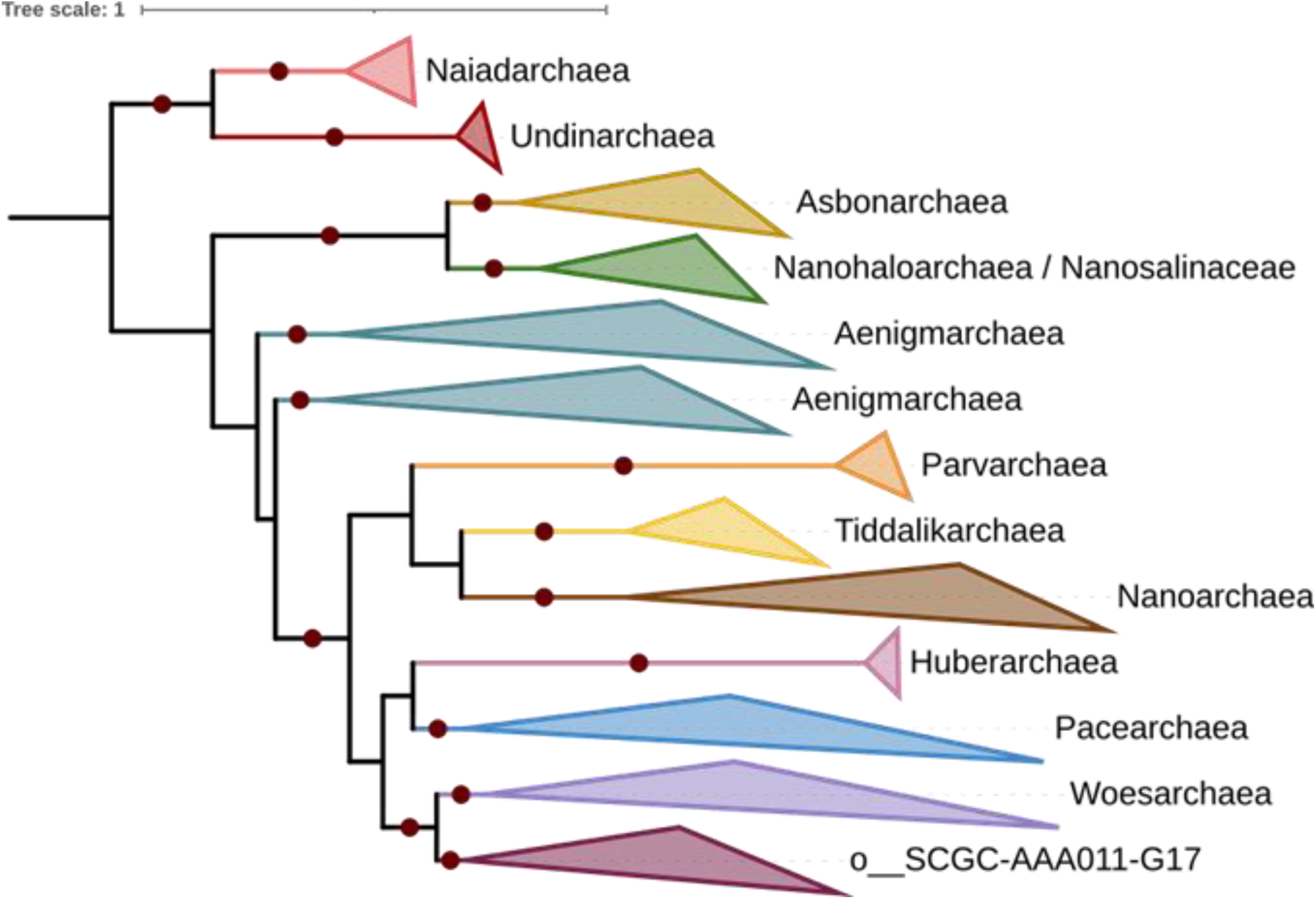
Phylogeny of the Nanostetteria DNA primase. The tree has been rooted according to Dombrowski et al. 2019, using the PriS, PriL archaeal DNA primases as the outgroup.

We recovered in our phylogeny a clade grouping *Naiadarchaea* and *Undinarchaea*, well separated from DPANN cluster II, the sisterhood of *Asbonarchaea* and *Nanohaloarchaea/Nanosalinaceae*, and the close relationships between *Nanohaloarchaea* and *Aenigmarchaea* (Figure 8). The latter were divided in two clades located between *Nanohaloarchaea* and a large clade including *Nanoarchaeota, Huberarchaea, Parvarchaea, Woesearchaea, Pacearchaea* and the recently described *Tiddalikarchaea* (Vazquez-Campos et al., 2021). The *Nanostetteria* DNA primase tree obtained was therefore in good agreement with our insertion analysis, except for the RNA polymerase B insertion that suggests that *Aenigmarchaea* and *Nanohaloarchaea* are sister group (Figure 9). Our tree is also consistent with most recent DPANN phylogenies based on the concatenation of marker proteins (Dombrowski et al., 2019, 2020a,b, Aouad et al., 2022, Baker et al., 2024, 2025, Wen-Cong Huang, 2025). This indicates that the archaeal two subunit DNA primase (PriS, PriL) was replaced by the monomeric DNA primase in the branch of the archaeal tree leading to *Nanostetteria*. Our phylogenetic analysis thus refutes the LGT hypothesis and indicates that the monomeric primase present in a subgroup of DPANN can be considered as a synapomorphy testifying for the validity of the clade *Nanostetteria*.

**Figure 8:**
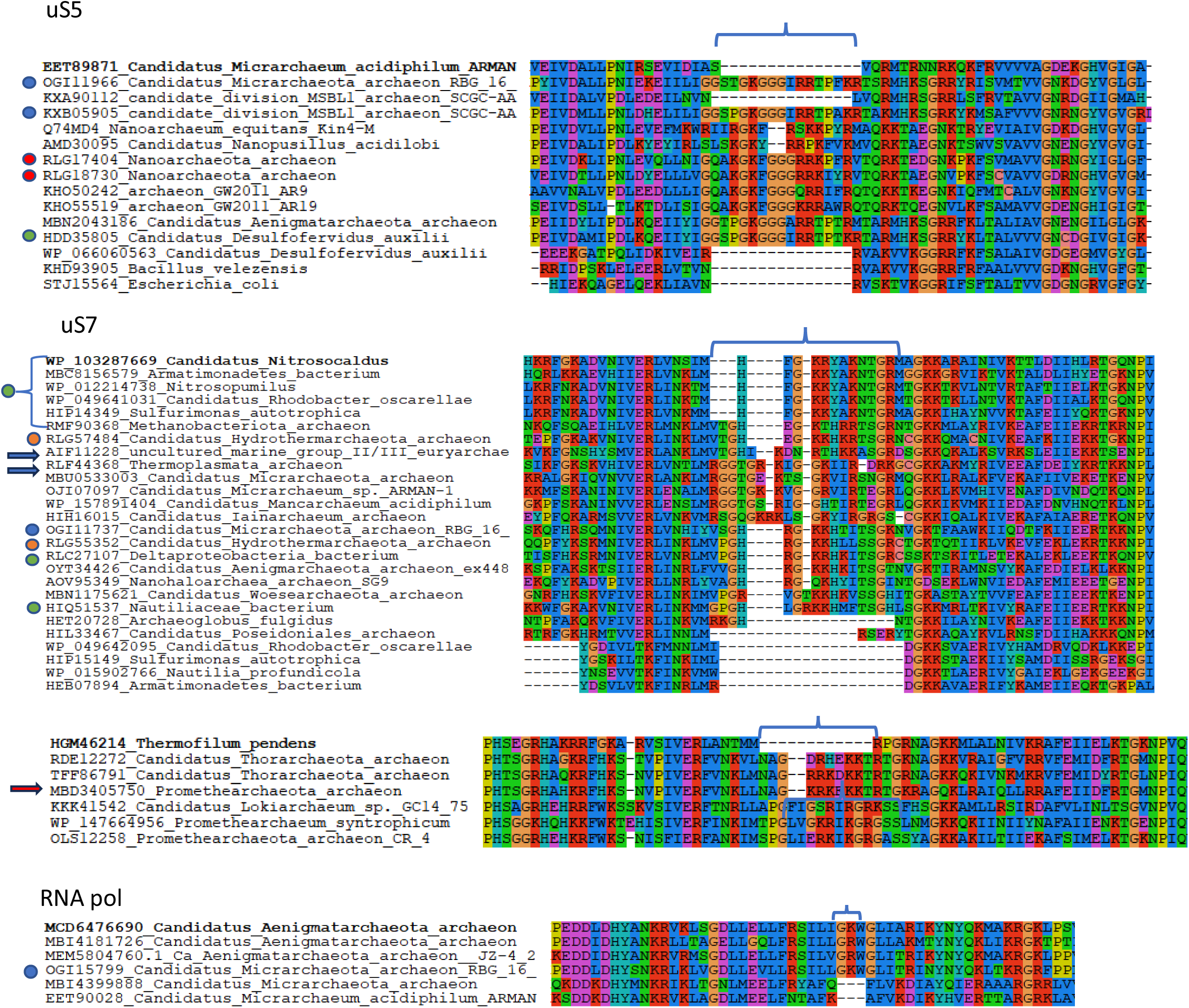
Detection of insertions in sequences of species whose relatives do not contain the insertion. Sequences corresponding to species whose relatives do not contain the insertion are labelled with dots or arrows (see the main text).

**Figure 9:**
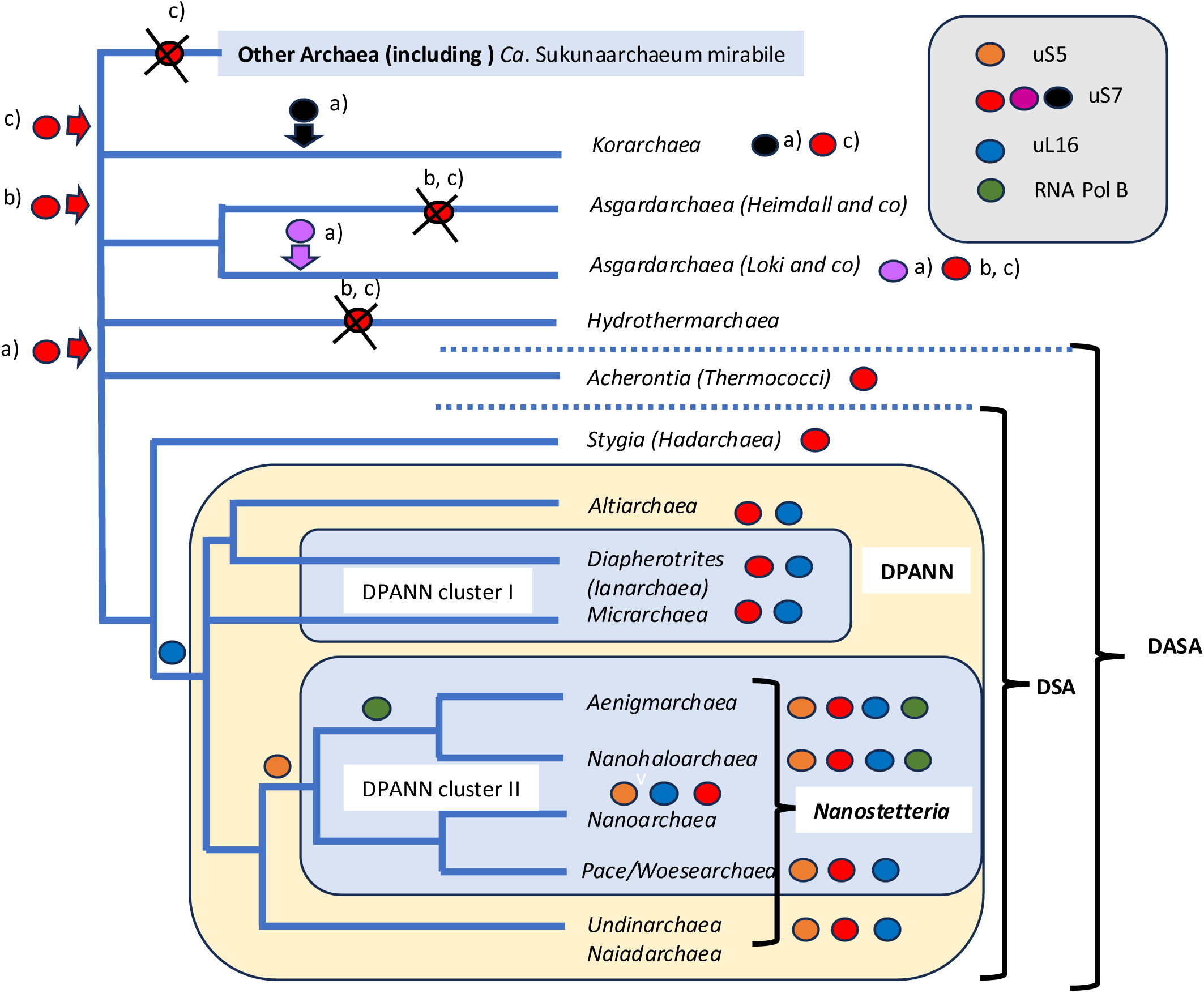
**Putative phylogenetic tree based on the different insertions detected in this work**. The figure illustrates the different clades proposed here based on the four different insertions (colored circles), red arrows (a) illustrate the hypothesis of three independent insertions for the uS7 insertion, (b) illustrate the hypothesis of a single insertion at the base of a clade grouping DASA and *Asgardarchaea* (c) illustrate the hypothesis of a single insertion at the base of a clade grouping DASA, *Korarchaea* and *Asgardarchaea*. The tree is unrooted because the exact position of the archaeal tree remains controversial (Forterre, 2025).

### Insertion analysis helps identifying MAG contaminations, misleading annotations and or lateral gene transfers

In several cases, we detected the insertions previously discussed in isolated MAGs supposed to belong to groups in which these insertions are normally not present. For instance, we identified the 15 amino acid uS5 insertion typical of *Nanostetteria* in the MAGs of one Micrarchaeon (RGB-16-49-10) and one Persephonearchaeon (candidate division MSBL1 archaeon) (blue dots in Figure 7, uS5). These insertions are not present in other MAGs of *Micrarchaea* or *Persephonarchaea*, suggesting that these two MAGs have either been (i) misannotated, (ii) contaminated by *Nanostetteria* sequences during genome reconstruction or (iii) the corresponding organisms recruited their uS5 gene from *Nanostetteria* by LGT. Notably, the putative proteome of Micrarchaeon RGB-16-49-10 also contains a uS7 protein with the 11 amino acids insertions typical of *Nanostetteria* and an RNA polymerase B” subunit containing the insertion of 3 amino acids detected in the RNA polymerase B” subunit of *Nanohaloarchaea* and *Aenigmarchaea* (blue dots in Figure 8, uS7 and RNA pol). One can conclude that this MAG has been misannotated and corresponds either to an Aenigmarchaeon or a closely related Nanostetterion.

We also detected the 15 amino acid insertions of uS5 in 18 MAGs annotated as “Nanoarchaeota archaeon” (two examples with red dots), whereas all sequences from isolated species of *Nanoarchaea,* such as *N. equitans*, contain instead a 13 amino acid insertion (two examples in Figure 8). This suggests that these “Nanoarchaeota” should correspond to a group of *Nanostetteria* distinct from the clade including isolated species of *Nanoarchaea*.

We also identified several other probable misannotations in analyzing the uS7 insertions. The uS7 11 amino acids insertion typical of *Nanostetteria* is present in the two MAGs of *Hydrothermarchaea* (orange dots in uS7-A Figure 7). However, these MAGs lack the two other insertions typical of *Nanostetteria* as well as the monomeric DPANN primase. Moreover, *Hydrothermarchaea* branch outside of DPANN in phylogenetic analyses (Jungbluth et al., 2017, Carr et al., 2019, Rinke et al., 2021) suggesting that these MAGs are contaminated with *Nanostetteria* contigs.

The uS7 11-16 amino acids insertions typical of the DSA clade is also present in a MAG annotated as *Thermoplasmata* and in several MAGs annotated as *Euryarchaea* group II or group II/III (two examples with arrows in Figure 7). These are clear cases of mis annotation or LGT. Some MAGs annotated as *Euryarchaeota* encode uS7 protein with unique insertions that can be nevertheless aligned with those of DSA members, suggesting that these MAGs correspond to new archaeal groups. Looking at the *Asgardarchaea*, we noticed a MAG annotated as “Promethearchaeota archaeon” that align with the inserton typical of *Thorarchaea* (Figure 8). All these observations suggest that misannotation (or lack of meaningful annotation) is a frequent issue in MAG analysis and that insertion analysis can be a valuable method to identify them.

In analyzing the uS5 and uS7 insertions, we surprisingly detected several bacterial MAGs encoding protein containing the archaeal insertions (see a few examples corresponding to green dots in Figure 8). We detected the uS5 insertion typical of *Nanostetteria* in two bacterial MAGs (see as an example the case of *Desulfofervidus auxilia* in Figure 8), four bacterial MAGs encoding uS7 containing the insertion typical of *Thaumarchaea* (not shown) and two containing an insertion that matches with the insertion present in uS7 of *Nanostetteria*: a Delta proteobacterium and a *Nautiliaceae* bacterium (Figure 8). Significantly, these insertions are not present in other MAGs or in *bona fide* genomes from *Bacteria* of the same groups. This indicates that bacterial MAGs encoding these proteins are in fact archaeal MAGs or were contaminated with archaeal DNA. During this work, we find indeed many cases of archaeal sequences annotated as *Ca*. bacterium. For instance, using as query in a BLASTP search an archaeal sequence (*Methanocella*) of the RNA polymerase region containing the 3 amino acids shared by *Nanohaloarchaea* and *Aenigmarchaeota*, we found 74 archaeal RNA polymerases present in bacterial MAG clusters (28/02/2026).

## Discussion

In this work, we identified four significant insertions in universally conserved proteins (three ribosomal proteins and the B” subunit of RNA polymerase) that clearly group together *Nanohaloarchaea* with DPANN archaea, as sister group to *Aenigmarchaea* and members of DPANN cluster II (*sensu* Dombrowski et al., 2020). This indicates that adaptation to high salt environment have taken place independently in *Halobacteria* and in *Nanohaloarchaea*, as previously suggested (Narasingarao et al., 2012, Baker et al., 2024). We failed to detect insertions grouping *Nanohaloarchaea* with *Haloarchaea* or with *Methanocellales* in the 37 universal markers that we analyzed. Our observations thus question the results based on protein markers in which *Nanohaloarchaea* were sister group to *Haloarchaea* (Petitjean et al., 2014, Rangel et al., 2021, Feng et al., 2021) or to *Methanocellales* (Aouad et al., 2019). The sisterhood between *Nanohaloarchaea* and *Haloarchaea* observed in some analyses could be explained by LGT of some protein markers between *Nanohaloarchaea* and their Haloarchaeal hosts. Moreover, in looking at the alignments of the 37 universal proteins used in our analysis, we notice the presence of a large number of indels in haloarchaeal proteins (Table S1). It is thus possible that some of them exhibit a fast evolving character that could attract fast evolving nanohaloarchaeal proteins. In the case of the sisterhood of *Nanohaloarchaea* and *Methanocellales*, this result was obtained either using recoding strategies or removing fast evolving positions (Aouad et al., 2019). This could explain the unexpected outcome of these analyses since amino acid recoding or the removal of fast evolving positions can sometimes produce misleading topologies by removing valid phylogenetic signal (Hernandez and Ryan, 2021, Rangel and Fournier, 2023).

Two of the insertions studied here (uS7 and uL16) support the specific affiliation of *Undinarchaea* and *Naiadarchaea* with DPANN cluster II. This suggests that these two lineages form a clade that is sister group to DPANN cluster II, as previously observed in phylogenetic analysis (Dombrowski et al., 2020). We suggest here calling *Nanostetteria* the robust clade grouping DPANN cluster II, *Undinarchaea* and *Naiadarchaea*. In agreement with this proposal, *Nanostetteria* are the only *Archaea* sharing the unique monomeric DNA primase that evolved from the fusion of the classical archaeal PriS and PriL primase subunits (Raymann et al., 2014, Dombrowski et al., 2020). The presence of this specific DNA primase can thus be considered as a synapomorphy of the clade *Nanostetteria*.

Notably, ranking this clade is not possible with present ICSP nomenclatures. In the traditional nomenclature, this rank would be located between a phylum and a superphylum, whereas in the ICSP nomenclature, this clade (being a subgroup of the kingdom *Nanobdellati*) would correspond to a taxon with intermediate position between the phylum and the kigdom levels. However, such level of taxa is not recognize in the ICSP nomenclature. In our opinion, this problem well illustrates the bias created in the ranking procedure by the fast-evolving character of DPANN.

Comparison between the phylogenies of this DNA primase (Figure 7) and the DPANN phylogeny recently published (Dombrowski et al., 2019, 2020, Baker et al;, 2024) indicates that this DNA primase co-evolved vertically after its introduction in an ancestor of *Nanostetteria*. The *Nanostetteria* primase branch far from the PriS and PriL primase subunits of DPANN cluster I in a global phylogeny obtained from the alignment of regions homologous between PriS and PriL with the corresponding regions of the *Nanostetteria* DNA primase (Dombrowski et al., 2020, figure S59). This indicates that this primase did not originated from a fusion that take place during the evolution of DPANN, but more likely from the monomeric DNA primase of a mobile element that infected an ancestor of *Nanostetteria* and replaced the ancestral two subunits archaeal primase still present in DPANN-*Archaea* cluster I and in *Altiarchaea*.

In recent phylogenies, *Altiarchaea* appeared to be sister group to DPANN-*Archaea* (Kellner et al., 2018, Rinke et al., 2021, Castelle et al., 2021, Aouad et al., 2022) or sister group to a clade grouping *Diapherotrites* and *Micrarchaea* (Dombrowski et al., 2019). In the latter case, *Altiarchaea* could be considered themselves as members of DPANN-*Archaea*. Our analysis confirms the close relationships between *Nanostetteria*, DPANN cluster I and *Altiarchaea*, since they share clearly homologous insertions in uS7 and uL16. The similariry between the insertion of the *Altiarchaea* uS7 and those of *Dipherotrites* suggests a possible clade grouping *Altiarchaea* and some lineages of DPANN cluster I, in agreement with the result obtained by Dombrowski and colleagues. We could not find insertion specifically grouping *Micrarchaea* and *Diapherotrites* with *Nanostetteria* at the exclusion of *Altiarchaeales*. Our observations thus only support the monophyly of DPANN-Archaea if *Altiarchaea* are included in this superphylum.

The uS7 insertion indicates a close relationships between DPANN (including *Altiarchaea*) and *Stygia* (*Hadarchaea*) suggesting the existence of a clade grouping all these lineages (here called the DSA clade). Finally, the large uS7 insertion suggests the possibility of an even larger clade grouping DSA lineages with *Acherontia* (*Thermococcales, Methanofastidiosa, Teinoarchaea*) (here called the DASA clade) which has also eluded previous phylogenetic analyses (Figure 9). The grouping of DPANN with *Thermococcales* was previously recovered by ancient phylogenetic analyses in which *N. equitans* and *Parvarchaea* turned out to be sister group to *Thermococcales* (Brochier et al., 2005, Brochier-Armanet et al., 2011).

Notably, the new clades proposed here include several archaeal lineages that still have the phylum status in the ICSP nomenclature, and thus should be considered as super phylum (for DPANN) or even “super, super phylum” (for DSA and DASA) clearly indicating that present nomenclature should be reconsidered.

In their recent phylogenetic analysis, Baker and colleagues have located DPANN (including *Altiarchaea*) as sister clade to group II *Euryarchaea* (*sensu* Forterre et al., 2025, a clade grouping *Thermoplasmatoa* and *Halobacteriota* in the ICSP nomenclature). We don’t find insertions supporting this topology. In fact, three deletions of the large uS7 insertions would be required to reconcile the presence/absence of this insertions with the published topology of Baker and colleagues. The presence of large uS7 insertions shared at the same position by all DASA members and several other archaeal lineages (*Thaumarchaea*, *Asgardarchaea* and *Korarchaea*) is better explained by three independent acquisitions of these insertions, as suggested by their lack of significant sequence similarity (hypothesis *a*, red, black and purple arrows in Figure 9).

In the case of *Thaumarchaea*, the independent acquisition hypothesis is supported by the lack of this insertion in closely related lineages (*Bathyarchaea, Aigarchaea*). The situation is less clear in the case of *Asgardarchaea* and *Korarchaea* since the position of these lineages is less well established in archaeal phylogenies and they are sometimes located rather close to DPANN-*Archaea* (Baker et al., 2024). The monophyly of *Asgardarchaea* is strongly supported by phylogenetic analyses and by the presence of a unique protein, such as the asgardactin, in all members of the group (Zaremba-Niedzwiedzka et al., 2017, Da Cunha et al., 2017, 2019, Liu et al., 2021, Aouad et al., 2022, Rodrigues-Oliveira et al., 2022, Eme et al., 2023). It is thus likely that the uS7 insertions in DASA and *Asgardarchaea* have taken place independently (purple and red arrows in Figure 9). Alternatively, if these insertions are homologous (hypothesis *b,* red arrow in Figure 9), they possibly originated in an ancestor of a large clade grouping *Asgardarchaea* and DASA. In that case, one should imagine that they were later lost in the branch leading to the clade grouping all lineages of *Asgardarchaea* lacking this insertion and that rapid sequence divergence explains why these insertions now exhibit very low sequence similarity. In that scenario, one can even imagine that the insertion present in the uS7 protein of *Korarchaea* is homologous to those previously discussed (hypothesis *c,* red arrow in Figure 9), and originated from a very ancient insertion that was lost in both *Crenarchaea* and some *Asgardarchaea*.

Interestingly, we failed to detect the insertions studied here, in particular the uS7 one, in *Hydrothermarchea* and in *Ca*. Sukunaarchaeum mirabile. *Hydrothermarchea* are usually located close or within DSAG and *Asgardarchaea* at the base or sister group to *Euryarchaea* (Carr et al., 2019, Rinke et al., 2021, Baker et al., 2025). The absence of the DSAG insertions in *Hydrothermarchaea* support their localization outside of this group (Figure 9). The recently described *Ca*. Sukunaarchaeum mirabile has the smallest archaeal genome reported to date and is obviously an obligatory parasite, which has been found associated to a dinoflagellate (Harada et al., 2025). The proteins of this miniature archaeon evolve rapidly and it exhibits an extremely long branch in an archaeal phylogenetic tree, precluding up to to determine its phylogenetic position (Harada et al., 2025). Despite the ressemblance in size and lifestyle with DPANN-Archaea, it lacks the uS7 and uL16 insertions that we found in all DPANN-Archaea (Harada, personal communication) indicating that it most likely does not belong to this superphylum.

Several studies have shown that indels can contains significant phylogenetic information that is often neglected (Tule et al. 2024; Redelings et al. 2024). This issue is particularly relevant in the case of highly conserved ribosomal proteins, where substitution rates are low due to their organisation in a huge protein and RNA complexe, in such case indel events may represent an important complementary signal. The analyses reported here suggest that systematically looking for insertions in proteins conserved between all *Archaea* and/or between *Archaea* and *Eukarya* could indeed help determining the actual history of the domain *Archaea*, confirming previous clades suggested by phylogenetic analyses, as in the case of the specific insertion that we detect in *Acherontia* (Figure 7), but also reaveling some unexpected new ones and their connections. This methodology could be of course also used to tackle difficult problems in bacterial and eukaryotic phylogenies.

Beside the four insertions detected here, we noticed many other insertions and deletions (indel) in the 36 universal DPANN-Archaea proteins that we analyzed. These indels usually took place in regions conserved in *Archaea* and are often variable within DPANN-*Archaea*. Generally speaking, the alignment of DPANN-*Archaea* protein sequences with those of their archaeal homologues clearly require to introduce more indels than the alignment of most proteins from other archaeal lineages with their archaeal homologues. This indicates that their proteins evolve much more rapidly than most other archaeal proteins, which has probably help DPANN-*Archaea* to adapt to their unique host-dependent lifestyle. Remarkably, the insertions detected in ribosomal proteins have taken place in regions that are close to RNA sites important for translation (Figure 3), suggesting that they could reduce the efficiency and fidelity of protein synthesis in DPANN-*Archaea*. It would be interesting to test this hypothesis by studying the effect of such insertions *in vivo* or *in vitro* on translation efficiency and fidelity.

The fast evolving character of DPANN-*Archaea* proteins explains why they are difficult to localize in the archaeal tree. This confirms that DPANN-*Archaea* should not be used in phylogenetic analyses whose objectives is to reconstruct the topology of the tree of life (Da Cunha et al., 2017, 2022) or to determine the topology of the archaeal tree. The fast evolving character of DPANN-*Archaea* also explains why these *Archaea* are still divided in several phyla in the ICSP nomenclature. In the future, it will be especially important to consider these differences in evolutionary rates in the ranking and classification procedures. Our observations suggests that the relative number of insertions in conserved proteins such as universal ones could be an interesting method to determine more accurately the relative rate of protein evolution in order to determine the actual evolutionary distance between lineages in ranking-based nomenclature.

Finally, a byproduct of our analysis was the detection of insertions in proteins from groups in which these insertions are normally absent. Some of these anomalies could be due to LGT, but we suspect that most of them correspond to contaminations leading to misannotation of these proteins. A striking examples of misannotations is the presence in the NCBI database of archaeal proteins that are annotated as bacterial proteins. Beside providing new insights on the phylogeny of organisms, well defined insertions can thus be very useful to rapidly identify misleading annotation, and possibly help the correct binning of MAGs, once programs to take into account this potentially new tool will be available.

## Materials and methods

### Screening and analysis of insertions in subsets of universal proteins

We started our analysis by screening for insertion/deletions (indels) the alignment of the 36 universal proteins used by Spang and colleagues (Spang et al., 2015) to determine the position of *Asgardarchaea* in the tree of life. During this analysis, we noticed that the sequences of universal proteins from *N. equitans* exhibit many unique insertions and deletions (indels) (see an example in Figure S38 of Da Cunha *et al*. 2017), in agreement with the idea that these episymbionts with reduced genome are fast-evolving organism (Brochier *et al*., 2005). To prevent the problems created in phylogenetic analyses by fast evolving species, we had previously removed DPANN-*Archaea*, *Korarchaea*, *M. kandleri* and sequences from unknown species from the species dataset of Spang and colleagues and produced 36 new untrimmed alignments (Da Cunha *et al*. 2017). We then added back *N. equitans* sequences and noticed that 38 indels were necessary to align the 36 universal proteins of *N. equitans* analyzed with those of other *Archaea*. Notably, the number of indels required to align other subgroups of *Archaea* with other Archaea was much lower, varying from 1 for many individual species up to 17 for *Haloarchaea,* except for the three Asgard available at that time (two *Lokiarchaea* and one Hodarchaeon) since 50 indels were required to align their sequences between themselves and with non *Asgard* archaeal species (Table S1).

We then added seven DPANN, including two *Nanohaloarchaeota* (*Nanosalinarium*, *Nanosalina*), two sequences from *Altiarchaea*, and 11 sequences of *Asgard* archaea our 36 alignments. We were interested to compare the extent of insertions in DPANN-*Archaea* and *Asgardarchaea* that both use to live in symbiosis/association with other organisms (Dombrowski et al., 2020, Imachi et al., 2020).

Using the same species dataset, we also made a hand-made alignment of the ribosomal protein uS5 (where u stands for universal according to the new system for naming ribosomal proteins, Ban et al., 2014) that was not present in the initial markers, but that one of us (PF) originally use to test the possible relationships between *Lokiarchaeota* and *Eukaryotes* (unpublished observation).

We obtained new untrimmed alignments, and we searched by eyes for indels in conserved regions of these proteins, focusing on *Nanohaloarchaea*. As in the case of *N. equitans*, we noticed that universal proteins of *Nanohaloarchaea* exhibit more indels than most other archaeal lineages, testifying for their fast-evolving character. Many of these indels have occurred in variable regions (especially in the N and C-terminal regions) and are difficult to interpret. Some of them are not conserved between different members of the same DPANN lineage, either testifying again for the fast-evolving character of these organisms and/or suggesting problems in metagenome reconstruction. However, we could identify four insertions that have taken place in regions otherwise very well conserved in the three domains or at least between *Archaea* and *Eukarya*? Three of these insertions were present in the ribosomal proteins uS5, uS7 and uL16 and a fourth one in the B” subunit of the DNA-dependent RNA polymerase. We then analyzed in more detail the four insertions, adding by hand sequences from other relevant archaeal groups and performing careful hand-made alignments. In some cases, we detected these insertions in one or a few MAGs from an archaeal group in which the vast majority of sequences do not contain these insertions (see some examples in Figure 8). BLAST searches using these odd sequences always revealed that they correspond in fact to other groups containing the corresponding insertions. We suspected that these exceptions could be due to MAG contamination or rare LGT and we discussed some of them in a separate section. We also noticed a 6 amino acid deletion present in the ribosomal proteins uL22 of *Nanohaloarchaea* and all DPANN in a region otherwise strictly conserved in size in the three domains of life. However, we don’t analyze this deletion because the homology between different deletions cannot be tested via amino acids sequence similarity.

We performed hand-made alignments of the regions corresponding to the four insertions detected here (available on request). To prepare the figures, final alignment using subsets of sequences were obtained using BLOSUM30 matrix with MAFFT and iterative refinement methods L-INS-I or G-INS-I (Katho et al., 2019). The alignments were visualized using Seaview (Gouy et al., 2021. Structure-based alignment of some universal protein was performed using the ESPRIPT web-server 3.0 (https://espript.ibcp.fr/ESPript/ESPript/) (Robert and Gouet, 2014).

### Phylogenetic analysis

For the DNA primase phylogenetic analysis, we chose 154 primases sequences from representatives of the diversity of *Nanostetteria*. Protein alignment was done with MAFFT V7 with the amino acid matrix BLOSUM 30 and iterative refinement methods L-INS-I or G-INS-I (Katoh et al., 2019), the same strategy was used for the alignment of universal proteins. The trimming was performed with BMGE with a BLOSUM30 matrix and the -b 1 parameter (Crisculo and Gribaldo, 2010) after trimming, the matrix contains 409 positions. The Maximum likelihood trees were constructed using the IQ-TREE multicore version 2.1.4 software (http://www.iqtree.org/) (Nguyen et al., 2015) with the ModelFinder option -MFP, to the best substitution model that was LG+F+R7. The branch robustness was estimated by the SH-aLRT approximate likelihood ratio test (Guidon et al., 2010) the ultrafast bootstrap approximation (Hoang et al., 2028) (10,000 replicates for each).

---

**Figure S1.**
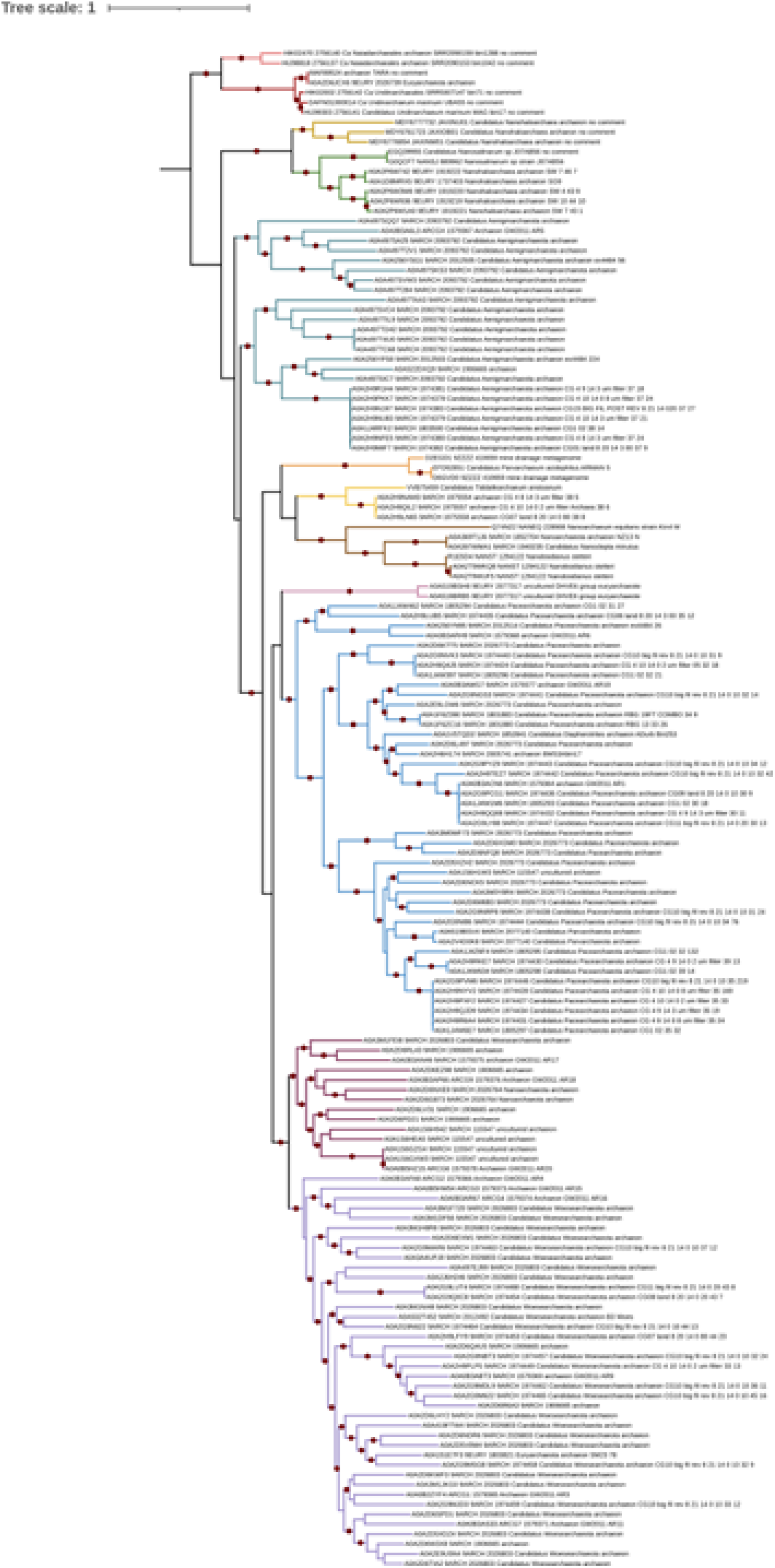

